# PCPpred: Prediction of Chemically Modified Peptide Permeability Across Multiple Assays for Oral Delivery

**DOI:** 10.64898/2026.01.19.700485

**Authors:** Akshay Shendre, Pushpendra Singh Gahlot, Gajendra PS Raghava

**Affiliations:** Department of Computational Biology, Indraprastha Institute of Information Technology, Delhi-110020, India

**Author notes:** **Mailing Address of Authors** Akshay Shendre Pushpendra Singh Gahlot Gajendra P. S. Raghava (GPSR). **<u>Corresponding Author</u>** Prof. Gajendra P. S. Raghava, Head and Professor, Department of Computational Biology, Indraprastha Institute of Information Technology, Delhi, Okhla Phase 3, New Delhi, India – 110020, Website: http://webs.iiitd.edu.in/raghava/.

**Keywords:** Modified peptides, Cyclic peptides, Permeability, Descriptors, Physicochemical properties, Molecular Fingerprints, Machine learning, Stacked ensemble, Chemical language models

## Abstract

Chemically modified peptides, including cyclic peptides, have emerged as promising candidates for oral delivery yet they face the challenge of low membrane permeability. In this study, the datasets were sourced from CycPeptMPDB, a database for membrane permeability of peptides obtained from different assays. Our quantitative analysis showed a clear discordance between permeability measured using PAMPA and cell-based assays (Caco-2, MDCK, and, RRCK), thereby explaining its limits as surrogate for cell-based assays. Therefore, we developed assay-specific predictive models to more accurately capture permeability determinants in each system. We systematically compute diverse features of modified peptides using open-source software and used fine-tuned peptide embeddings generated using pretrained chemical language models. Baseline models were developed using the generated multi-hierarchical molecular features. We also developed a stacked ensemble architecture, which utilizes multi-hierarchical features in models as base learners. The ensemble model achieved the best PAMPA test set performance with an MSE of 0.200, R^2^ of 0.685, and PCC of 0.830; and a R^2^ of 0.783 on Caco-2 test set. Model trained on 2D Mordred descriptors attained the highest performance on the Caco-2 test-set with MSE of 0.129, R^2^ of 0.793, and PCC of 0.892, surpassing state-of-the-art approaches such as CPMP. To support widespread adoption, we developed an open-access web-server (https://webs.iiitd.edu.in/raghava/pcppred/) for users to design modified peptides using human comprehensible MAP (Modifications and Annotations of Proteins) format, converting MAP to SMILES format, and predict permeability across assays with result visualization. To ensure widespread adoption, and reproducibility, we also provided a standalone on GitHub (https://github.com/raghavagps/pcppred).

## 1. Introduction

Peptides with cyclization represent a promising and increasingly vital category of bioactive molecules due to their unique conformational rigidity [1], enhanced resistance to enzymatic degradation [2], and ability to engage challenging targets [3]. Unlike linear peptides, these peptides maintain stable tertiary structures, enhanced target binding affinity and pharmacokinetic stability. Additionally, the cyclization constrains the backbone and thereby facilitates the formation of intramolecular hydrogen bonds. These bonds which can shield the polar groups therefore facilitate the passive membrane diffusion, which is a key requirement for oral bioavailability and intracellular activity [4,5]. Mechanistically, the permeability of these membrane active peptides is governed by a delicate balance between polar surface area, molecular size, and their conformational dynamics. Studies have shown that intramolecular hydrogen bonding and solvent-exposed polar residues critically influence transcellular transport. For instance, Rezai et al. (2006) in their study demonstrated the adoption of membrane permeable conformations by certain cyclic scaffolds which can temporarily mask polar groups, enabling efficient diffusion across lipid bilayers [5]. As a result of recent discoveries, the clinical relevance of peptides including the peptides with cyclization continues to grow. Several drugs including, cyclosporine (an immunosuppressant), Zilucoplan (a complement C5 inhibitor for myasthenia gravis), and Afamelanotide (used in erythropoietic protoporphyria) reflect the broad utility of the peptides. The recent FDA’s approvals of several macrocyclic peptides such as Rezafungin further reflects the trend of developing peptides as therapeutic agents, especially in antimicrobial and autoimmune domains. A 2022 review by Ramadhani et al., reflects the potential of cyclic peptides acting as anticancer agents [6]. Further reviews including Chia et al 2023., emphasized the expanding use of cyclic peptides as antiviral agents, in infectious diseases [7], highlighting their high selectivity and bioactivity as reasons for continued investment in this class.

Consequently, the study and optimization of peptide properties, particularly their permeability, is essential not only for advancement of peptide-based therapeutics but also to enable standard design frameworks in drug discovery pipelines. However, despite the therapeutic advantages, peptides with cyclization often face challenges in drug development, primarily due to poor membrane permeability, thereby limiting the oral bioavailability, and limited ability to reach intracellular targets [8,9]. Peptide’s membrane permeability is primarily influenced by physicochemical properties, including hydrogen bond (donor/acceptor) counts, polar surface area and molecular weight [10,11]. The solvent-exposed hydrogen bond donors in lipophilic environments critically impact permeability, as peptides with numerous exposed amide NH groups tend to exhibit reduced lipophilicity and passive permeability [8,11,12].

Experimental assays such as Parallel Artificial Membrane Permeability Assay (PAMPA) [13], Caco-2 permeability testing [14], Ralph Russ Canine Kidney (RRCK) [15] and Madin-Darby Canine Kidney (MDCK) cell assays [16] remain the gold standard for evaluating peptide permeability, however these assays are resource-intensive, and time-consuming, thereby not suited for large-scale screening. This limitation led to an ever-growing demand and interest to utilize computational approaches to predict physicochemical properties including the permeability at early stages of drug development. Several recent studies including CycPeptMP, Multi_CycGT, PeptideCLM, CyclePermea, PharmPapp and CPMP have made significant advancement to this field by utilizing deep learning, graph neural networks and transformer based modeling and prediction approaches [17-22]. However, these studies often depend on limited or features derived using commercial softwares, difficult to replicate or fail to offer public tools. These limitations hinder widespread adoption and reproducibility, particularly in academic and resource constrained settings. Thereby to address these limitations, we propose a machine learning framework which includes stacked ensemble machine learning based model architecture that integrates predictions from multiple base learners trained on distinct features derived from SMILES representations [23] of the peptides. These multi-hierarchical features include, atomic features derived using RDKit [24], 2D and 3D molecular descriptors obtained via Mordred [25], PaDEL [26], and RDKit, molecular fingerprints from PaDELPy (a python wrapper for PaDEL-Descriptor software), and RDKit; and SMILES embeddings fine-tuned from pre-trained chemical language models. Each of the base learner (e.g., random forests, SVMs, GBMs) produces permeability predictions that are further used as features by machine learning regressors (e.g., XGBoost), creating a robust and generalizable predictive model architecture. Additionally, for widespread adoption of the computational approach of permeability prediction, a web server has also been developed to democratize public access to our models. This platform allows users to design peptides in MAP (Modifications and Annotations of proteins) [27] format which is a human readable format under development by our group. The design module of the server can be used to first design the desired peptide. The designed peptide can then directly be used as input to our prediction tool to predict permeability of peptide through the prediction module of the server. The prediction module of the web server, if given the input in MAP notation first converts it to SMILES notation then automatically computes relevant features, subsequently runs the trained models to predict permeability for different permeability assays based on user input and finally allows visualization of the result. This tool hence empowers users from diverse backgrounds to rapidly evaluate novel membrane active peptides without proprietary software dependencies.

## 2. Materials and Method

### Datasets

In this study, peptide data from the CycPeptMPDB database [28] was utilized, specifically employing experimentally obtained logarithmic membrane permeability values (LogP_exp) as the target variable. The CycPeptMPDB compiles permeability measurements obtained from a range of assay platforms, including PAMPA, Caco-2, MDCK, and RRCK permeability assay. The dataset comprises 7,298; 1332; 64; and 186 entries corresponding to PAMPA, Caco-2, MDCK, and RRCK assays in the latest available version (v1.2) of the database. To construct the working datasets for each assay type, the datasets were curated, by first removing the unreliable/inaccurate entries, (entries with assay Permeability value <= -10) then, by removing the duplicate entries (entries with same SMILES representation and molecular weight). To remove duplicates, the average of the permeability values was taken, thereby taking a holistic representation of all the studies. These new entries representing the average permeabilities of the duplicate entries were given new Id’s, starting from the last entry present in the particular dataset. This resulted in a total of 6960, 1259, 64, 175 entries for PAMPA, Caco-2, MDCK, and RRCK assays. All the peptides present in either of the assay types have either circular or lariat shape, and none of the cyclization were because of the disulfide bonds between two cysteine residues. In PAMPA, out of 6960 peptides 4660 peptides were circular whereas 2300 peptides have lariat shape. Similarly, for Caco-2, 658 peptides have circular and 601 have lariat shape. RRCK dataset have 175 peptides with only circular shape and MDCK have 55 circular and 9 lariat shape peptides. The datasets were subsequently partitioned into training and testing subsets. As the permeability of peptides are influenced by physicochemical parameters, including molecular weight, initially, the entries were sorted in descending order based on molecular weight. Figure S1, shows the dispersion of molecular weights of these peptides for each dataset. To ensure stratification across the molecular weight spectrum, every fifth datapoint from the sorted dataset was assigned to the Test set, resulting in an approximate 80:20 split between Train and Test sets. This sampling approach was employed to maintain a uniform representation across the chemical space of the peptides in the test set.

### Feature Extraction

To comprehensively characterize the modified peptides with cyclization and facilitate robust permeability prediction, a diverse array of molecular features from both their monomeric sequence and SMILES representations were extracted to capture characteristics across multiple hierarchical levels. To encode sequence-level information, CycPeptMPDB annotations were used to obtain Monomeric composition-based features. These features included natural analog based composition (represented by 21 dimensional vectors corresponding to 20 canonical amino acid residues and 1 unidentified non-natural residue), and monomer specific composition consisting of 385 dimensional vectors corresponding to distinct monomer types. Both the types of representation quantified the relative abundance of each residue within a particular peptide sequence, thereby enabling fine-grained learning of chemical diversity and variation within a sequence. In addition to that, atomic-level features computed using RDKit obtained elemental composition, bonding characteristics, and charge distribution through 23 features. These included the relative abundance of key atoms (Br, Cl, F, C, N, O, S), atomic degree counts, bond-type frequencies, formal charge, and indicators for aromaticity and ring topology. Together, the monomeric and atomic features provide a detailed encoding of peptide composition, bonding, and atomic level structure.

Structural, substructural, and contextual features were subsequently obtained to capture physicochemical and topological information from SMILES representations of peptides. Two and three - dimensional molecular descriptors were obtained using RDKit, Mordred, and PaDEL-descriptor. These open-source tools collectively yielded descriptors that quantify molecular geometry, topology, hydrophobicity, electronic distribution, among other properties which are key determinants of membrane permeability. Molecular fingerprints were generated using RDKit and PaDELPy to capture structural motifs and connectivity patterns. The fingerprints included Morgan and count-based Morgan fingerprints (2048 bits each), capturing circular atom environments using extended-connectivity fingerprint (ECFP) like representations [29]; and twelve additional fingerprint types such as EState [30], MACCS Keys [31], PubChem, Substructure, Klekota-Roth [32], and Atom Pair [33] fingerprints, representing both local atomic environment and global molecular framework. Finally, high-level representations were captured by fine-tuning pretrained transformer-based chemical language models including ChemBERTa-zinc-base-v1 [34], PubChem10M_SMILES_BPE_450k [35], and MoLFormer-XL [36], on peptide SMILES strings to obtain the embeddings. The embeddings were obtained from the final hidden layers and mean-pooled to 768 dimensional vectors, to capture contextual rich information by integrating chemical syntax and semantics.

### Baseline Models

To evaluate the predictive utility of various molecular feature representations for peptide permeability prediction, a suite of eleven machine learning regression algorithms (Light Gradient Boosting Machine (LGBM), Extra Trees, Random Forest, Gradient Boosting, AdaBoost, and XGBoost regressors, as well as individual regressors including Decision Tree, Linear Regression, Support Vector Regressor (SVR), K-Neighbors Regressor (KNN), and Multi-Layer Perceptron (MLP) Regressor) were implemented utilizing different features. Before model training, feature pre-processing was done by removing constant or non-numerical features, feature standardization using StandardScaler, and replacement of missing or Nan (not a number) with zero. Models were trained and validated using five-fold cross-validation for statistical robustness and reduced overfitting. Performance metrics used to evaluate models included Mean Squared Error (MSE), Root Mean Squared Error (RMSE), Mean Absolute Error (MAE), Coefficient of Determination (R^2^), Pearson Correlation Coefficient (PCC), and Spearman Correlation Coefficient (SCC).

Individual baseline models were constructed using features derived from different molecular representations described in section 2.2. For sequence level modeling, the suite of machine learning models based on natural analog and monomer specific compositions were used to evaluate the predictive capacity of these features under identical pre-processing and validation procedures. Similarly, atomic-level features describing elemental characteristics were modeled independently as well as combined with monomer specific composition features, to assess their predictive capability and contribution towards predicting permeability. Structural descriptors consisting of 2D and 3D molecular features, extracted from SMILES were used to develop separate and combined descriptor-based models, thereby exploring the influence of topology, geometry and physicochemical properties on permeability prediction. Further, to capture the influence and utility of substructural motifs, molecular fingerprints of 14 types were used individually and combined together to train the suite of machine learning models. Furthermore, embeddings derived from fine-tuned chemical language models were employed as high-level chemical syntax and semantic representation to train the suite of machine learning models. Finally, machine learning models integrating multiple feature types (atomic, structural (2D/3D descriptors), fingerprints, and language model embeddings) were developed to utilize their complementary strength after pre-processing the features. The pre-processing of features in models integrating multiple feature types involved, removal of low-variance features (variance < 0.05), and eliminating highly correlated features (|r|>0.9) for individual feature types then concatenating the features. These concatenated features were further undergone feature selection to remove redundancy. This was achieved by eliminating the feature which have lower correlation with permeability if features are highly collinear (|r|>0.9). The models were then trained based on top n features (n ∈ 10, 20, 50, 100, 200, 500) derived based on feature importance score calculated using random forest regressor. Collectively, these baseline models were trained to prepare an evaluation framework to benchmark the contribution of individuals and their combinations towards permeability modeling.

### Chemical language Models

In addition to SMILES derived features, pretrained chemical language models (mentioned in 2.2 section) were used to directly learn structure-property relationships from SMILES representations of the peptides. The model checkpoints were fine-tuned, on the training datasets. Each SMILES sequence was tokenized using the associated pretrained tokenizer with a maximum input length of 325 tokens. Input sequences were padded or truncated as needed to maintain uniformity. A regression head was appended to the pretrained model, and the network was fine-tuned end-to-end using a MSE loss function. Training was performed over 20 epochs using the Adam optimizer with a learning rate of 5e-5 and a batch size of 16. The fine-tuned models were evaluated on test sets, and predictive performance was assessed using standard regression metrics mentioned in section 2.3.

### Stacked Ensemble Model Architecture

A robust two-stage stacked ensemble regression architecture that integrates heterogeneous molecular features and multiple machine learning algorithms was developed to predict peptide permeability. The overall framework consists of a pre-processing pipeline followed by a two-tiered learning strategy. The complete architecture is illustrated in Figure 1. Four distinct molecular feature sets were used as inputs to the model architecture which includes, 2D and 3D molecular descriptors, capturing a wide range of physicochemical and geometric properties; molecular fingerprints, representing binary patterns of substructures within the molecules; SMILES-based molecular embeddings, obtained from a fine-tuned MoLFormer-XL chemical language model that encodes chemical syntax and semantics; and atomic features, encoding atomic composition and elemental properties.

**Figure 1.**
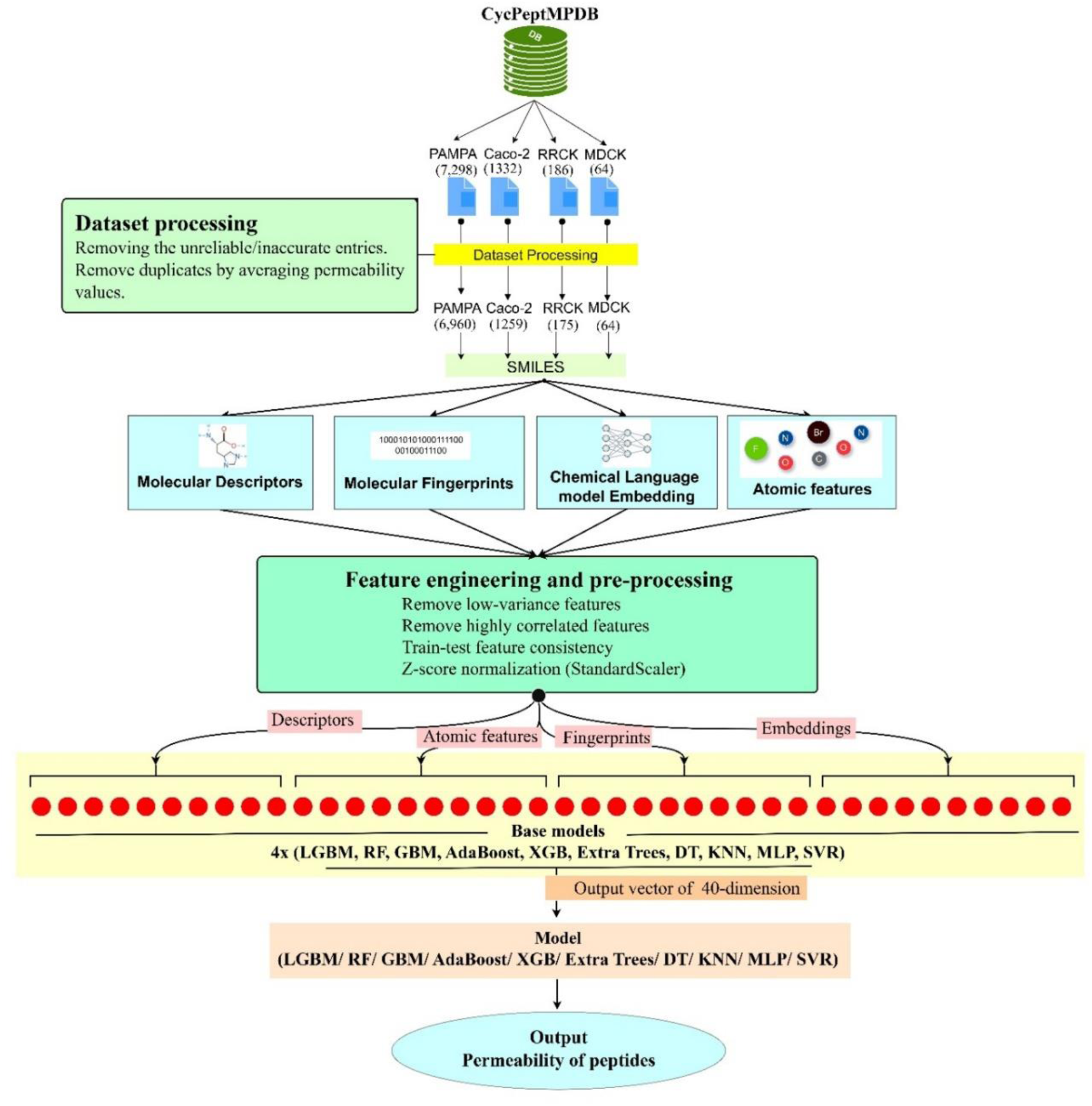
Schematic overview of the stacked ensemble machine learning architecture to predict membrane permeability of peptides. The schematic shows the complete pipeline including data pre-processing, feature generation and predictive modeling using stacked ensemble architecture.

Each representation was subjected to a consistent pre-processing pipeline, which involved steps involving, removal of constant and low-variance features (variance < 0.005) to eliminate noise and redundancy; correlation-based feature selection, where highly collinear features (|r| > 0.9) were discarded unless strongly associated with the permeability value; train-test feature alignment, ensuring that only features preserved after pre-processing in the training set were retained in the corresponding test set and Z-score normalization (StandardScaler) to ensure numerical comparability across features and to accelerate convergence in optimization-based models. The model architecture consisted of two stages. Stage 1 consisted of base learner ensemble where for each pre-processed feature set, a panel of ten baseline regressors (LGBM, Random Forest, Gradient Boosting, AdaBoost, XGBoost, Extra Trees, SVR, KNN, MLP, and Decision Tree) were independently trained. A K-fold cross-validation strategy was used to train each base learner, with K=5. Out-of-fold predictions for each learner on the training set were stored as meta-features, while predictions on the test set were averaged across folds. The process was repeated separately for all four feature types, resulting in four meta-feature matrices for both training and testing. The final feature matrices were obtained by concatenating the predictions from all base learners across the different feature representations, yielding a unified and rich set of inputs for meta-modelling. Stage 2 of the architecture consisted of machine learning regressors, where aggregated features were used as input to the regressors to make permeability predictions.

## 3. Results

CycPeptMPDB, a cyclic peptide database utilized in this study compiled permeability measurements from assays including PAMPA, Caco-2, MDCK, and RRCK permeability assay. Processing of the datasets (removing inaccurate entries and averaging duplicates) resulted in a total of 6960, 1259, 64 and 175 entries corresponding to each assay type. PAMPA set consisted of 14 peptides containing only natural amino acid residues (L-amino acids), whereas 5635 of the peptides consist of at least one D-amino acid residue, and majority (>99%) of peptides consist of non-natural amino acid residues. Caco-2 dataset consists 9 peptides with only canonical L-stereoisomers of amino acids, 421 peptides have at least one D-enantiomeric residue and like PAMPA set, apart from 9 peptides with only natural residues, the rest have residues with modification or are of non-natural origin. Similarly, MDCK and RRCK sets also have 1 peptide each with only canonical L-stereoisomers of amino acids, and 51 and 144 peptides with at least one D-type amino acid residue.

Investigating whether permeability prediction should rely on assay-specific dataset, or a unified dataset combining all four assay types, we investigated descriptor-permeability relationships across PAMPA, Caco-2, RRCK and MDCK. The top 5 most positively and negatively correlated 2D/3D descriptors with permeability values in Caco-2 dataset (1259 entries) were explored across the assays. The correlation coefficients are summarized in Table 1, and their assay-specific variations are illustrated in the line plot (Figure 2A). Positively correlated descriptors such as AtomTypeEState.252, ATSC4c, and ATSC.4, maintained a similar trend in RRCK and MDCK, whereas polarity, and topology related descriptors (TPSA, TopoPSA, and TDB9) showed consistent negative trends. PAMPA, on the other hand showed weak or inverse correlations, indicating its mechanistic discordance from cellular systems. Inter-assay correlation analysis of descriptor-permeability correlation values (Table S1) revealed strong coherence among Caco-2, RRCK and MDCK (r =0.96-0.98) whereas a negative correlation of these assays with PAMPA, underscores a distinct permeability behaviour. Further comparing shared peptides (n=483) among assays, confirmed assay-dependent variability in the experimental permeability values as evident by a sample of peptide given in Table S2, (SMILES corresponding to each ID in Table S2 is given in Table S3) and through heatmap (Figure 2B). Next, to test the feasibility of a unified predictive model irrespective of the assay type, datasets from all assays were combined and 2D/3D descriptors were used as features to train machine learning algorithms. Models trained solely on descriptors achieved moderate performance (R^2^ = 0.446, MSE = 0.362, PCC = 0.672), while adding a four-dimensional assay-indicative vector markedly improved results (R^2^ = 0.602, MSE = 0.260, PCC = 0.777; Table 2). Complying with the observations, despite high concordance among cell-based assays, the divergent behaviour of the PAMPA which comprises the most prominent dataset, and low predictive capability of a generalized model underscores the necessity of developing assay-specific predictive frameworks to predict permeability of modified peptides.

**Figure 2.**
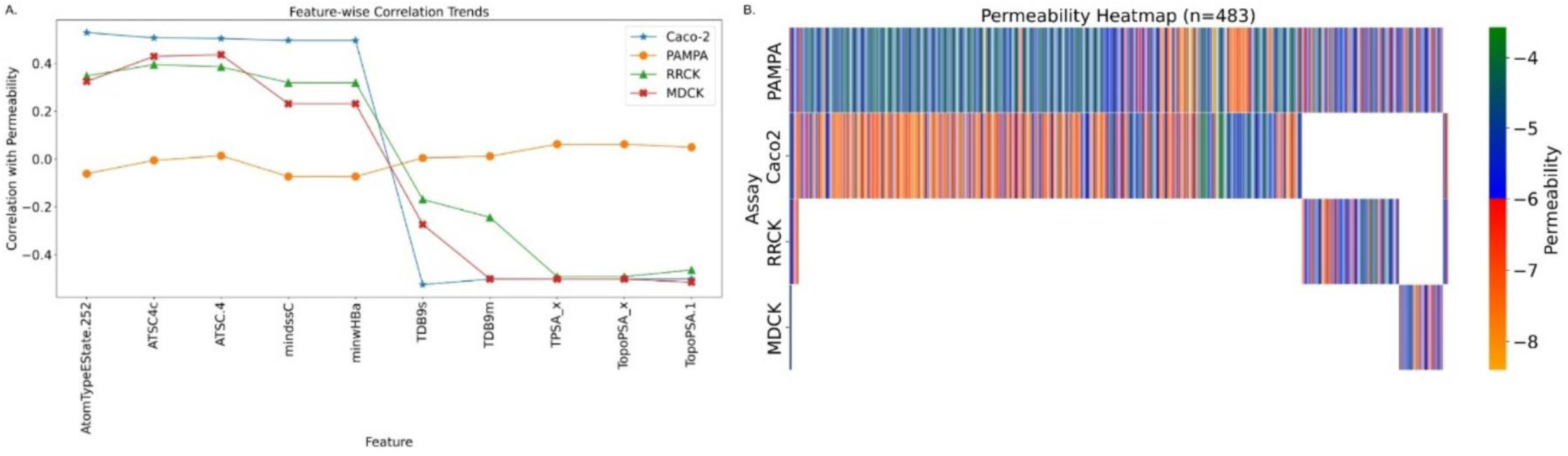
Inter-assay descriptor-permeability correlations and permeability values comparison of peptides. (A) Descriptor-permeability correlation trends of the top five positively and negatively associated descriptors across Caco-2, PAMPA, RRCK, and MDCK datasets. (B) Heatmap of permeability values for peptides, common among four assays (n=483), illustrating assay-specific variability and concordance.

**Table 1:**
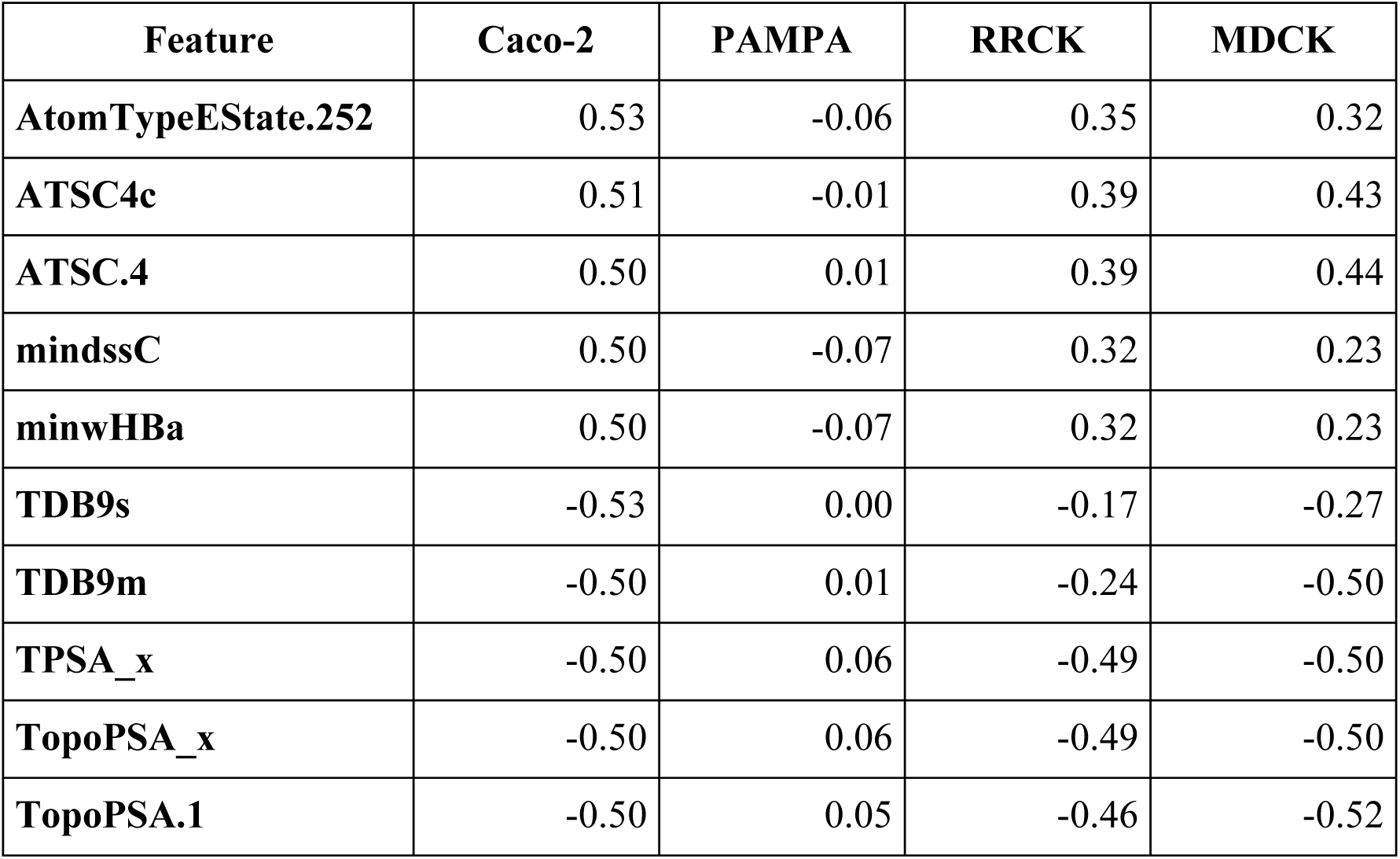
Correlation coefficients of the top five positively and negatively correlated 2D/3D molecular descriptors with permeability in the Caco-2 dataset, compared with their corresponding correlations in PAMPA, RRCK, and MDCK datasets.

**Table 2:**
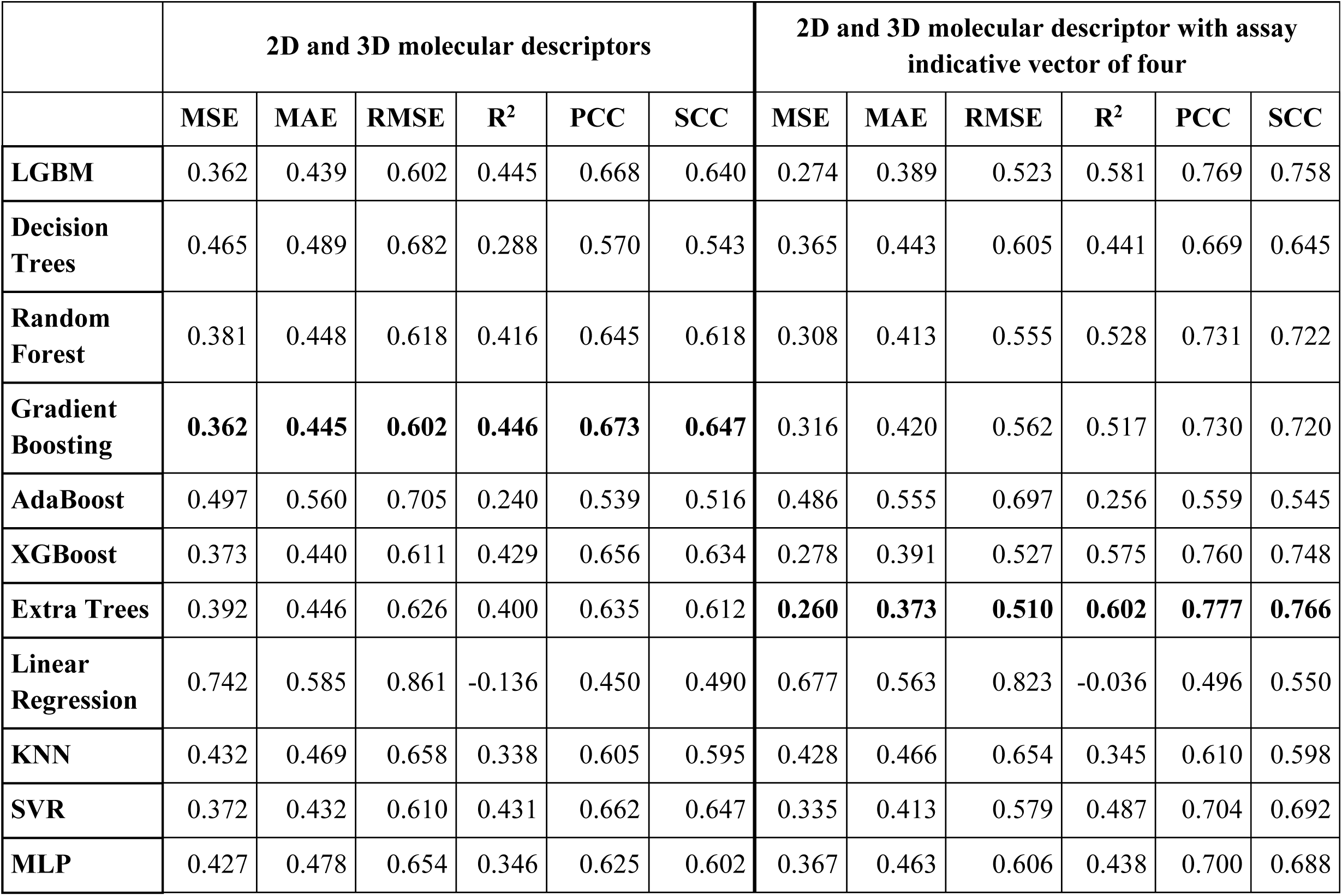
Test set performance comparison of generalized peptide permeability prediction models using 2D/3D descriptors alone versus descriptors with assay-indicative feature vector as features.

### Composition based analysis

Analysis of average amino acid composition obtained by mapping 385 monomers to their natural analogs (with ‘X’ representing undefined residues) revealed that peptides were rich in Leu, Ala, Phe, Pro, and Gly across all four assays, regardless of permeability classification (LogPexp >= -6.0 vs < -6.0) as indicated in Figure 3A, 3C, 3E, 3G. Examining full monomeric composition further indicated high frequencies of natural (Leu, Phe, Ala, and Pro), D-amino acid (dl, dP), and N-methylated residues (meL, meA, Me_dL, Me_dA) (Figure 3B, 3D, 3F, 3H). Next, to quantify the contribution of specific non-canonical modification, ten most frequent modified monomers in PAMPA and Caco2 sets were analysed for effect on permeability distributions. In PAMPA dataset, N-methylation and bulky hydrophobic residues such as N-methyl-leucine (meL), N-methyl-alanine (meA), N-methyl-D-leucine (Me_dL), N-methyl-D-alanine (Me_dA), were associated with higher permeability, shifting distribution above the -6 threshold (Figure S2). In contrast, D-proline (dP), D-leucine (dL), and N-benzyl-glycine (Bn_Gly) correlated with reduced mean permeability, consistent with a diffusion-driven mechanism where backbone N-methylation and hydrophobicity favour passive membrane permeation. In contrast, caco-2 showed a more complex, assay-specific effect (Figure S3). meL and meA reduced permeability, while presence of polar or conformationally restrictive modifications like Asp-piperidide and O-tert-butyl-serine [Ser(tBu)] enhanced permeability. D-residues (dL, dP) that were unfavourable in PAMPA improved permeability in Caco-2. These observations reflect the context dependence of structural modifications (N-methylation promoting passive diffusion in artificial membranes), cellular systems impose additional constraints such as solubility and efflux, allowing certain polar or sterically restrictive residues to enhance permeability under biological conditions.

**Figure 3.**
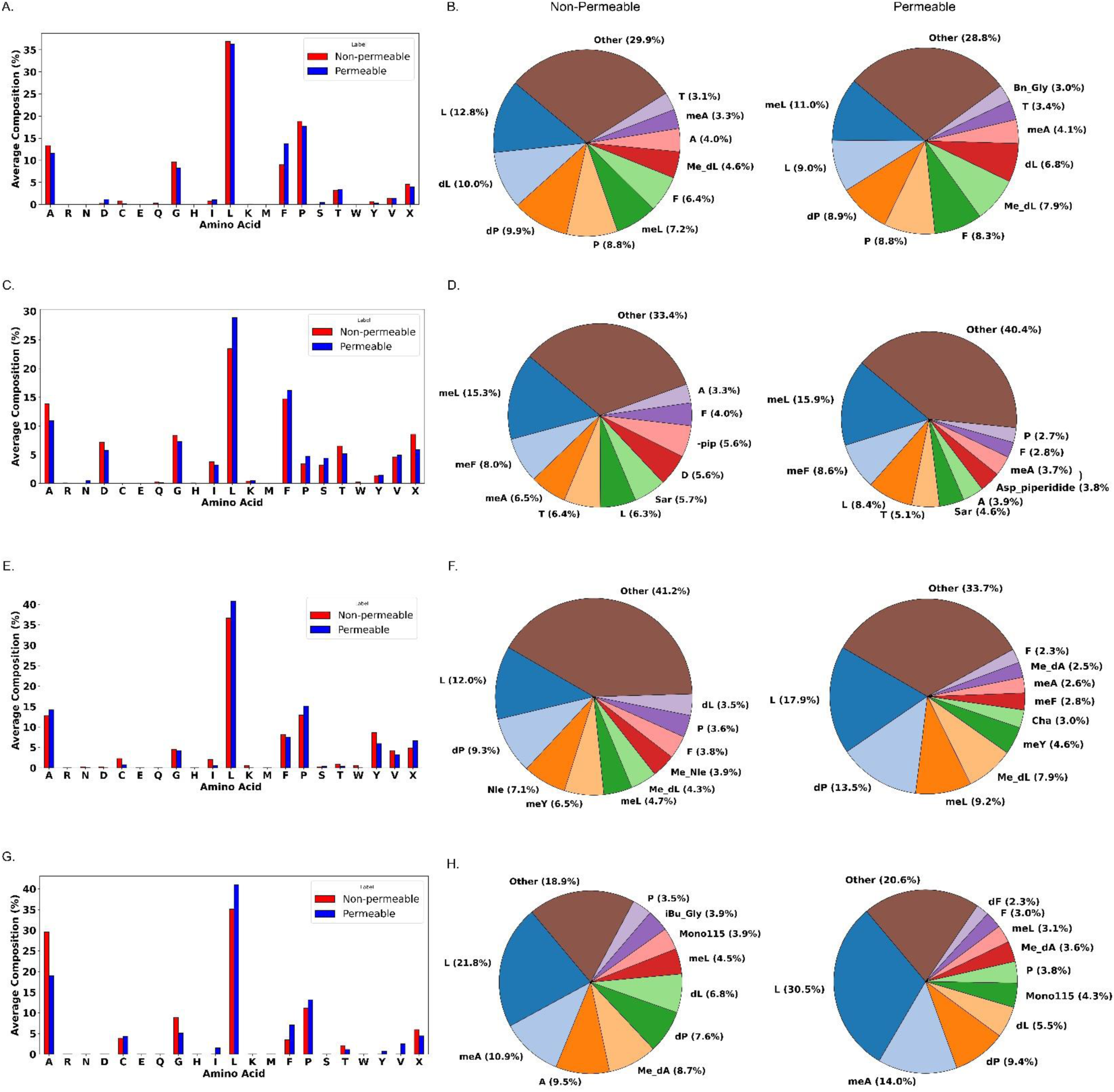
Amino acid and monomeric composition analysis of peptides across four permeability assay datasets. Bar plots shows the average amino acid composition (%) of peptides in the (A) PAMPA, (C) Caco-2, (E) RRCK, and (G) MDCK datasets. Blue bars represent permeable peptides and red bars represent non-permeable peptides. Pie charts representing the average monomeric composition (%) of the top 10 most frequent monomers in (B) PAMPA, (D) Caco-2, (F) RRCK, and (H) MDCK datasets. The left pie chart in each panel corresponds to non-permeable peptides and the right chart to permeable peptides.

### Molecular descriptors/physicochemical properties-based analysis

To extend our understanding beyond compositional factors, we analysed physicochemical properties and molecular descriptors governing permeability of peptides in PAMPA and Caco-2. Top 20 descriptors were chosen using feature importance scores from best-performing regressors (LGBM for PAMPA, Figure 4A; and Extra Trees for Caco-2, Figure 4D), and analysed for statistical significance, correlation trend, and class-dependent separation in box plots. In PAMPA, descriptors including MolLogP, minsOH, minssCH_2_, Chi.9, TopologicalCharge.12, MolecularId.7, and CrippenLogP emerged as influential candidates. MolLogP and CrippenLogP which are lipophilicity indicators showed positive correlation with permeability in non-permeable peptides but opposite correlation in permeable peptides, suggesting that excessive hydrophobicity may be hindering solubility or promote aggregation in train set (Figure 4B). Other structural indices like Chi.9, and minssCH_2_ showed an inverse correlation with permeable peptides in both train (Figure 4B) and test sets (Figure 4C), highlighting the role of topology and molecular branching in diffusion. Descriptors like minsOH, TopologicalCharge.12, and MolecularId.7 which are related to polar group exposure and charge distribution also showed a reversed trend between permeable and non-permeable peptides, implying that balanced electrostatic interactions and controlled polarity may support membrane transit by compensating role in balancing lipophilicity with solubility. The coherence of these patterns across train and test sets, along with statistically significant differences (Table S4) and distinct distributional differences evident from box plot analyses (Figure S4 and S5), reinforces their mechanistic relevance in artificial membrane diffusion.

**Figure 4.**
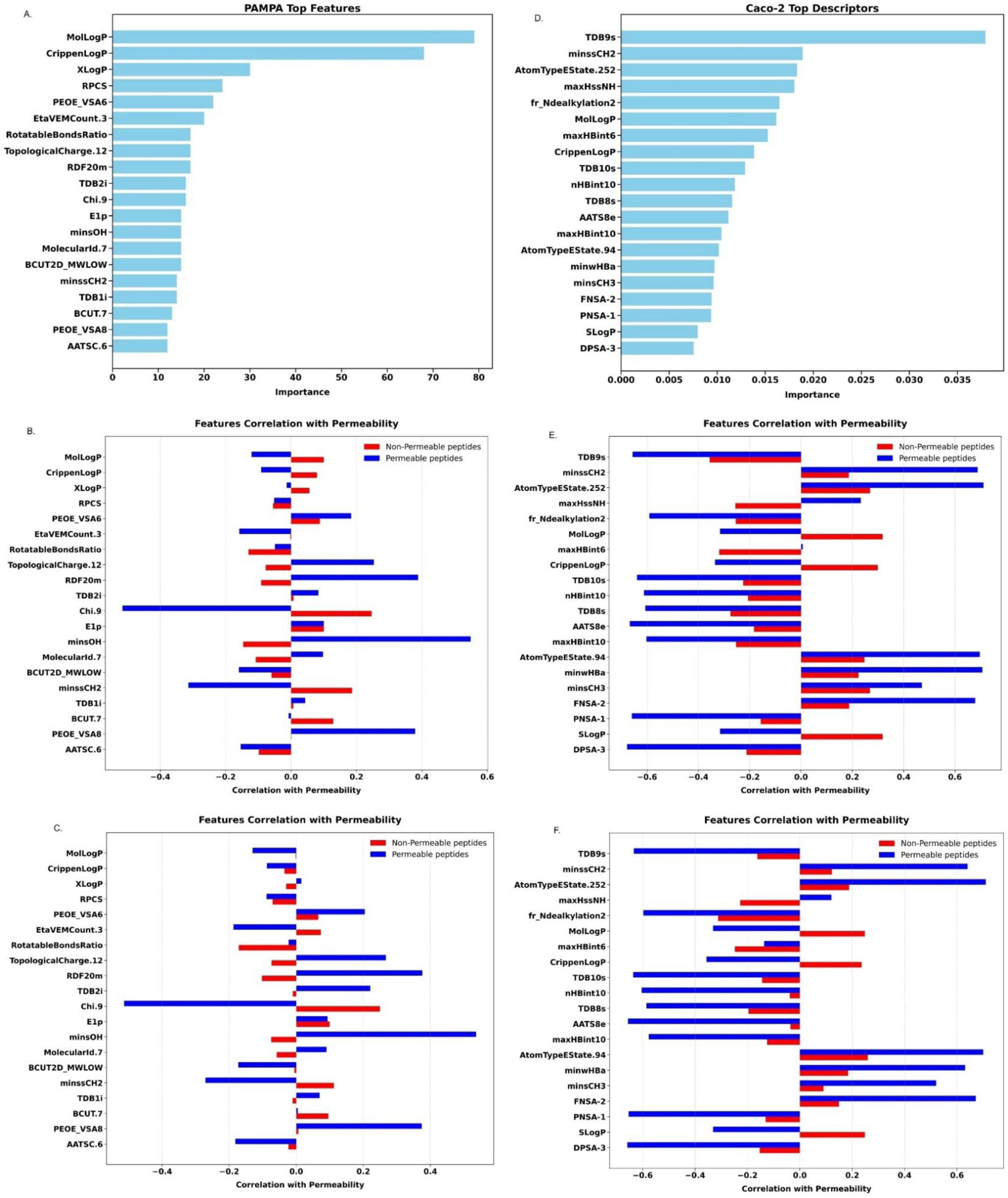
Top physicochemical descriptors influencing peptide permeability in PAMPA and Caco-2 assays. (A, D) Top 20 descriptors ranked by feature importance using LightGBM for PAMPA and Extra Trees for Caco-2. (B, C) Pearson correlation of key PAMPA descriptors with permeability, separated by permeable v/s non-permeable peptides in train and test sets, respectively. (E, F) Corresponding correlation trends for Caco-2 descriptors in train and test sets.

In Caco-2, descriptors like MolLogP, CrippenLogP, SLogP, maxHssNH showed consistent discriminatory effects between permeability classes (Tables S5). Similar to PAMPA, hydrophobicity related descriptors (MolLogP, CrippenLogP, and SLogP) showed positive correlations with permeability in non-permeable peptides but inverse correlations with permeable peptides (Figure 4E-4F), emphasizing towards excessive lipophilicity impeding cellular permeability, possibly through efflux interactions or limited solubility. In contrasts, maxHssNH, reflecting nitrogen-based hydrogen bonding capacity, exhibited opposite trends (positive correlation with permeability in permeable peptides, but negative in non-permeable peptides), indicating that conformational shielding or intramolecular hydrogen bonding may enhance diffusion by reducing polarity. These observations are further supported by distinct distribution in respective box plots (Figure S6-S7). Collectively, across both assays, descriptors corresponding to lipophilicity, charge distribution, and hydrogen bonding power reflected a repeated observation that moderate hydrophobicity and controlled polarity jointly governs permeability, whereas assay specific mechanisms like diffusion limits in PAMPA v/s efflux and solubility constraints in Caco-2 further modulate their influence.

### Permeability prediction of modified peptides with cyclization across assay types

In this section, we began by evaluating a diverse set of regressors using four different molecular representations, atomic-level features, monomeric composition, physicochemical descriptors, and molecular fingerprints. Each feature group was used to train multiple machine learning models on the datasets including PAMPA, Caco-2, RRCK and MDCK permeability dataset of peptides.

### Permeability modeling in PAMPA assay

We systematically evaluated the performance of baseline machine learning regressors models, chemical language models and proposed stacked ensemble architecture to predict the permeability of peptides derived from PAMPA assay. The train set consists of 5568 entries whereas the test set contains 1392 entries.

To evaluate the predictive utility of different molecular feature representations, we first trained regressors using atomic-level features, which quantify elemental composition, bonding patterns, and topological motifs through a 23-dimensional vector. These features attained a R^2^ of 0.554, and MSE of 0.283 (Table S6). Investigating monomer level composition features by representing each peptide using 385 distinct monomeric units and after removing invariant features yielded 243 features. This representation showed higher predictive performance compared to atomic features (MSE of 0.258, R^2^ of 0.595, and PCC of 0.772; Table S7). In contrast, simplification of monomers to their closest natural amino acid analogs (20 canonical residues plus ‘X’ for unknowns) markedly diminished prediction quality, with a highest MSE of 0.403, and reduced R^2^ of 0.367 and PCC of 0.607 as shown in Table S7. These findings suggest that while atomic descriptors capture general physicochemical patterns, detailed residue-specific information encoded at the monomer level plays a more critical role in governing passive permeability in modified peptides with cyclization. Importantly, combining both atomic and monomeric features (262 total features after removing invariant features) resulted in an improved overall performance achieving an MSE of 0.226, R^2^ of 0.645, and PCC of 0.803 (Table S6), indicating that the two representations together provide overall information gain and this synergistic improvement reflects the value of integrating traits from multiple hierarchies to capture diverse permeability-relevant attributes.

We further evaluated models trained on traditional molecular descriptors, including both 2D and 3D features derived from cheminformatics toolkits such as RDKit, Mordred, and PaDEL/PaDELPy. The modeling was undertaken to evaluate the utility of descriptor sets that encode fundamental topological, physicochemical, and three-dimensional structural properties. Such descriptors have widely been used in quantitative structure-activity relationship (QSAR) modeling and thereby provide a point of reference for evaluating the predictive utility in the context of cyclic peptide permeability. Models were trained on descriptors computed independently from each toolkit, and additionally on aggregated feature sets consisting of (i) all 2D descriptors, (ii) all 3D descriptors, and (iii) a combination of 2D and 3D descriptors. Among these, the regressor trained on Mordred 2D descriptors with 1227 features after removing constant features and 2D combined descriptors after removing constant features (2504), achieved the highest performances, with lowest MSE of 0.207, and R^2^ of 0.675. Notably, combining all 2d and 3D descriptors (2946 descriptors after removing invariants) yielded comparable performance (MSE 0.212, R^2^ 0.667, PCC 0.819), indicating that 2D descriptors by Mordred alone captures a wide spectrum of permeability-relevant properties as evident from Table 3.

**Table 3:**
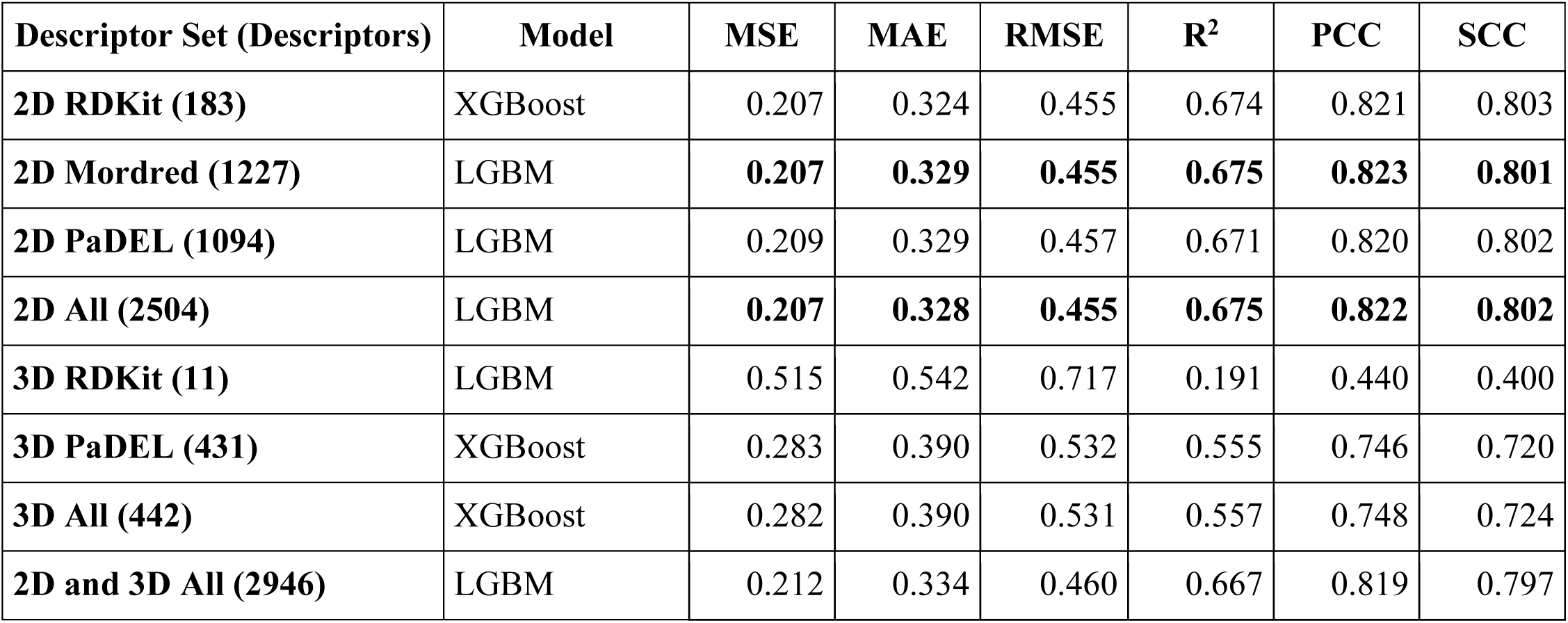
Best-performing regression models for each molecular descriptor set on the PAMPA dataset. The table reports test set performance across 2D, 3D, and combined descriptor sets using standard regression metrics: MSE, MAE, RMSE, R^2^, PCC, and SCC.

In the case of molecular fingerprints, which provide binary or count-based encodings of structural subcomponents, the best individual performance was observed using count based Morgan fingerprints (2048 bits), yielding an MSE of 0.226, R^2^ of 0.645, and PCC of 0.803 but the performance was further improved (MSE of 0.212, R^2^ of 0.666, and PCC of 0.816) when all the fingerprints types were aggregated together and invariant fingerprints were removed (6820 fingerprints), suggesting that despite their spare nature, fingerprints collectively encode meaningful substructural patterns relevant to membrane permeability (Table S8).

To assess the utility of contextual representations, chemical language models (CLM) were directly trained on smiles sequences of peptides. Three CLM’s were trained, of which are shown in Table S9. Among the fine-tuned models, PubChem10M_SMILES_BPE_450K achieved the highest performance, with an MSE of 0.281, R^2^ of 0.558. Although the performance was relatively inferior as compared to descriptor and fingerprint-based models, the results still promisingly demonstrate the capacity of sequence-based models to extract permeability relevant information from linear molecular notations. In a further extension, embedding vectors from these three pre-trained language models were extracted (768 vectors) and used as input to downstream regressors. Embeddings from MolFormer_XL_both-10pct language model achieved the highest modeling performance upon fine-tuning, yielding the test set performance, with a MSE of 0.206, R^2^ of 0.676, and PCC of 0.822. This notably surpasses other single modality models, suggesting that the embedding space learned by MolFormer_XL encodes rich structural and chemical patterns essential for accurate prediction of permeability (Table S10).

The complementary strengths of multi-hierarchical representations were leveraged further by combining features of modalities and selecting the top features based on feature importance using random forest. A model of top 50 features from all the modalities yielded the highest performance of MSE of 0.207 and R^2^ of 0.674 and PCC of 0.822 (Table S11). Second, we implemented stacked ensemble models architecture wherein base regressors were trained on different modality features (atomic features, molecular descriptors, fingerprints, and embeddings), and the predictions from each of the regressor were used to train meta-learner regressor. This modeling architecture achieved the highest overall performance, with an **MSE of 0.20, R^2^ of 0.685, and PCC of 0.830.** These results for stacked ensemble architecture are summarized in Table 4. These findings highlight the synergistic value of integrating diverse molecular representations through ensemble learning, underscoring its effectiveness in capturing complex structure activity relationships for improved predictive accuracy.

**Table 4:**
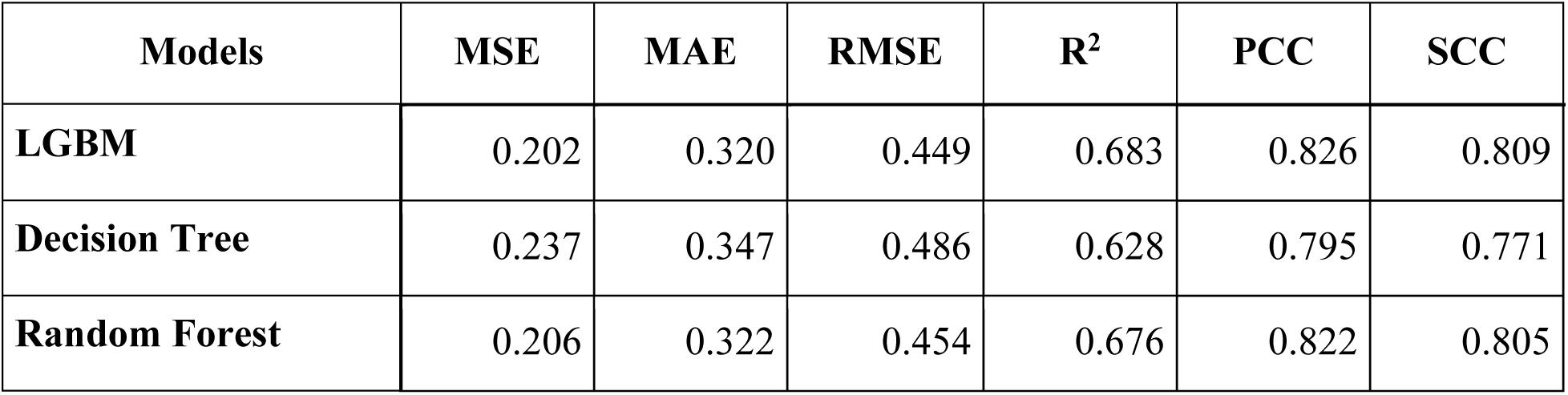
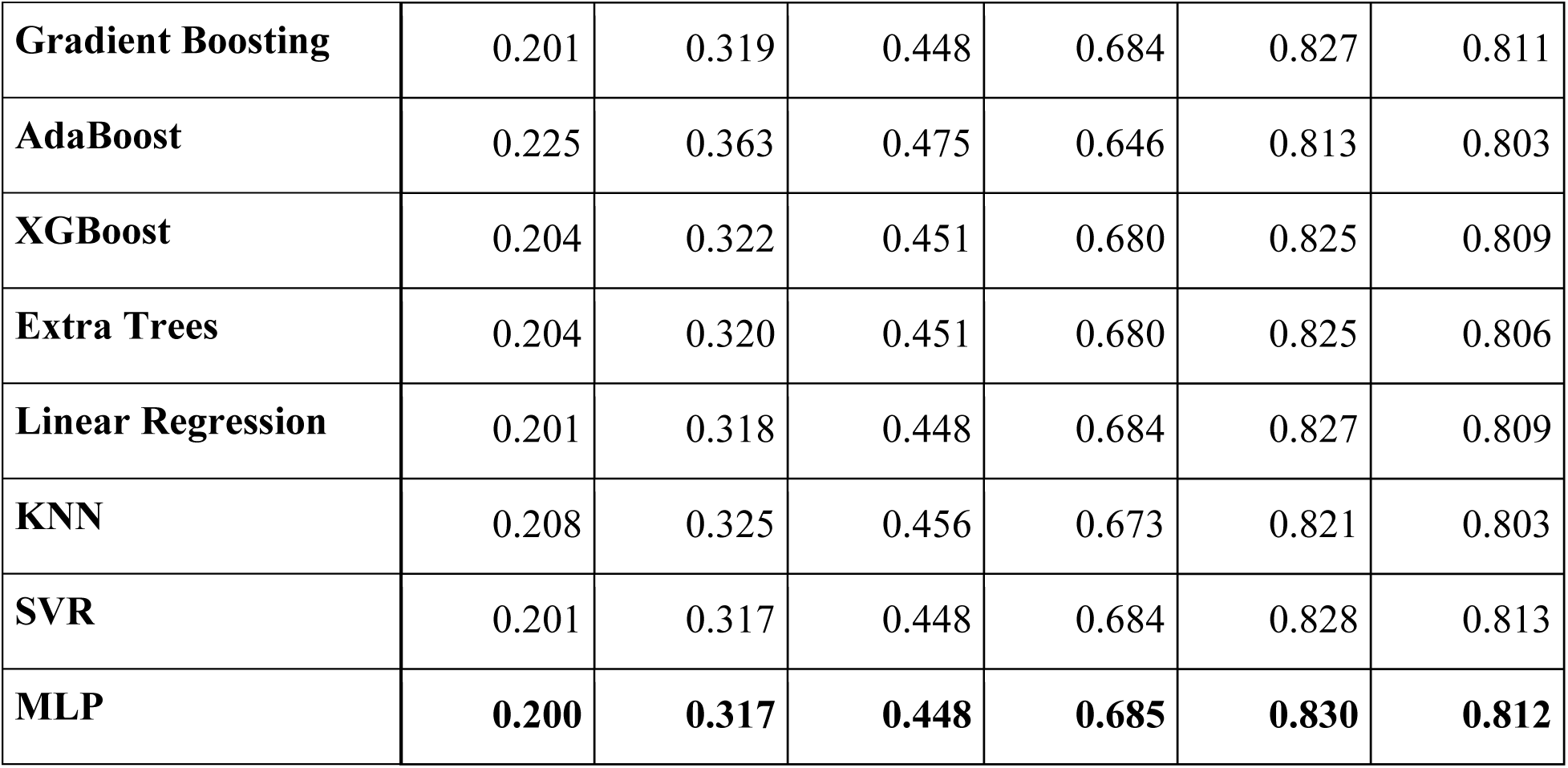
Test set performance metrics for stacked ensemble models for PAMPA assay.

To dissect the relative importance of individual representation types in the ensemble, an ablation study was conducted by systematically excluding base model trained on each feature type (atomic, descriptor, fingerprints, and embeddings) and evaluating the performance of stacked ensemble architecture using the MLP regressor as final model (highest performing model). The most pronounced performance degradation occurred upon exclusion of base models trained on descriptors evident by decrease in R^2^ in Figure 5A. Other metrics including MSE, MAE and RMSE (Figure S8) concurrently observed a significant increase upon exclusion of base learners trained with descriptors. The second most pronounced decline occurs when base learners trained on fingerprints are excluded, indicating their substantial role in capturing informative molecular patterns. In contrast, exclusion of embeddings leads to smaller, intermediate losses, while removal of atomic features has least impact. These results are shown in Figure 5A. The analysis thereby highlights that the final ensemble model, by effectively integrating diverse perspectives including topological, physicochemical, compositional, and learned sequence-level representations thereby achieved a robust and generalizable performance as summarized in Table 5.

**Figure 5.**
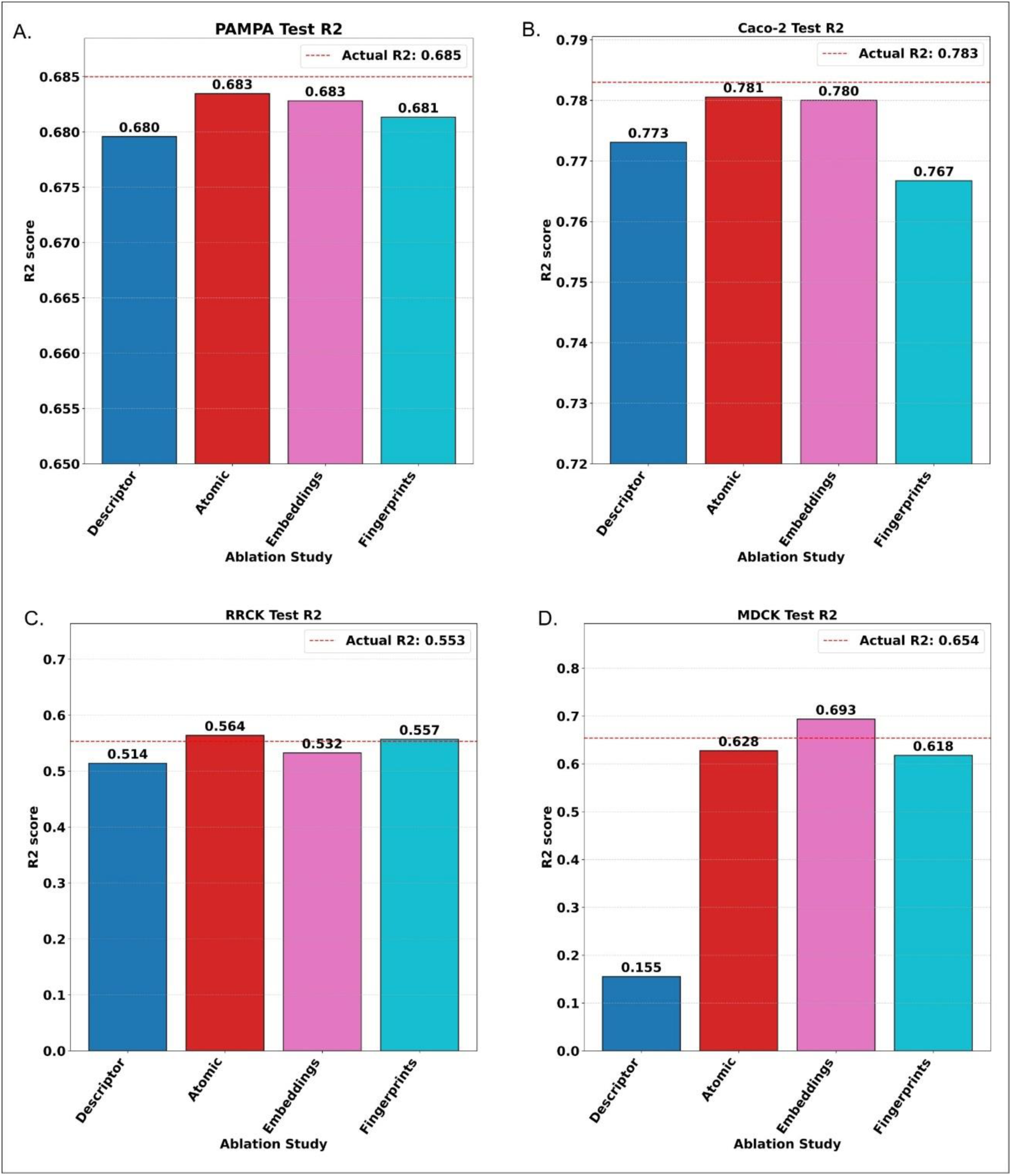
Ablation analysis of stacked ensemble architecture based on test set R^2^ performance across four permeability assays, (A) PAMPA, (B) Caco-2, (C) RRCK, and (D) MDCK. Each bar represents the model performance after excluding models trained on feature type, descriptors, atomic features, embeddings, or fingerprints from the stacked ensemble. The red dashed line represents the R² of the full architecture utilizing all four feature types, serving as the baseline for comparison.

**Table 5:**
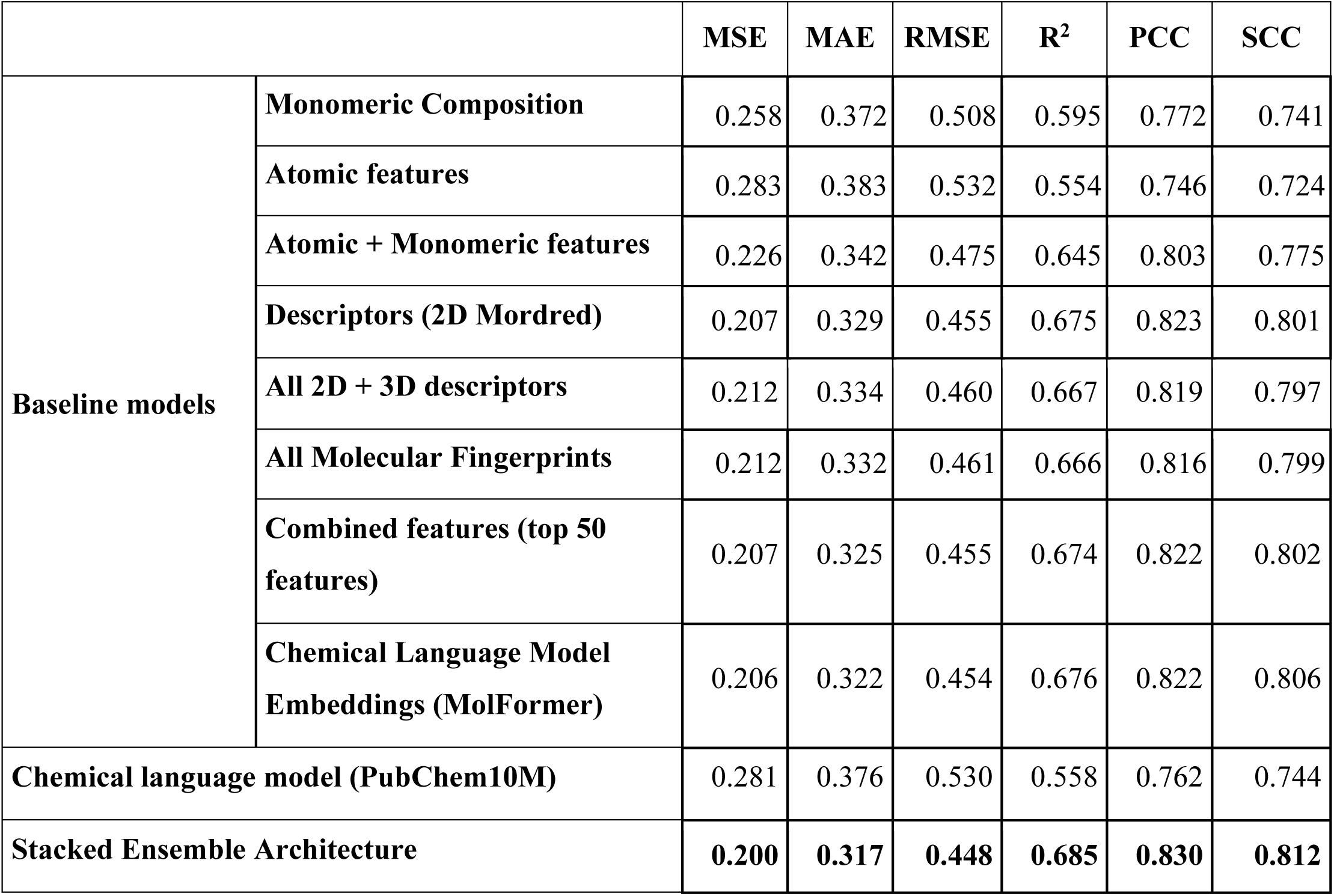
Test set performances of baseline models, chemical language model, and stacked ensemble architectures for PAMPA assay.

### Permeability modeling in Caco-2 assay

The same consistent modeling and performance evaluations approach was utilized to predict the permeability of peptides from the Caco-2 dataset as used in PAMPA assay. The train set consists of 1007 peptides whereas the test set consists of 252 entries. We employed the same regressor suite to train the baseline models. Initial evaluation of single-modality models revealed that atomic features (23 features), encoding the fundamental structural units and atomic connectivity patterns, outperformed composition based modeling wherein atomic feature based models achieved a R^2^ of 0.729 and a MSE of 0.169 (Table S12), in comparison to the monomer level composition (213 features after removing invariant features out of 385 monomeric feature vector) which shows lower R^2^ of 0.710 and a higher MSE of 0.181 in Table S13. However, similar to the observation in PAMPA assay, simplification of monomers to their closest natural amino acid analogs (20 canonical residues plus ‘X’ for unknowns) led to a further drop in performance (MSE of 0.229, R^2^ of 0.633), confirming the limited expressiveness of reduced alphabetic representations (Table S13). Further, combination of atomic and monomeric features (230 features after constant feature removal, Table S12) significantly improved performance (MSE = 0.138, R^2^ = 0.778, PCC = 0.883), establishing the synergistic effect of two types of features in membrane permeability.

In models trained on cheminformatics based physicochemical descriptor sets, it is observed that among all the models, regressors trained using 2D Mordred descriptors (1216 features after discarding invariant and non-numeric descriptors out of total 1616 descriptors) achieved the highest performance among all the models including stacked ensemble architecture with R^2^ of 0.793, PCC score of 0.892, and MSE of 0.129. Expanding to include all 2D and 3D descriptors resulted in reduced performance (MSE of 0.140, and R^2^ of 0.775). Table 6, highlights the complete test evaluation metrics for the best performers amongst all the different types of descriptors. Fingerprint based models showed reasonable performance, with Morgan count fingerprints (2048 bits) achieving MSE of 0.144 and R^2^ of 0.769, while aggregated fingerprint models (5503 fingerprints) achieved relatively higher performance (MSE of 0.134, R^2^ score of 0.785, and PCC of 0.887). The test set results for all other types of fingerprints are listed in Table S14.

**Table 6.**
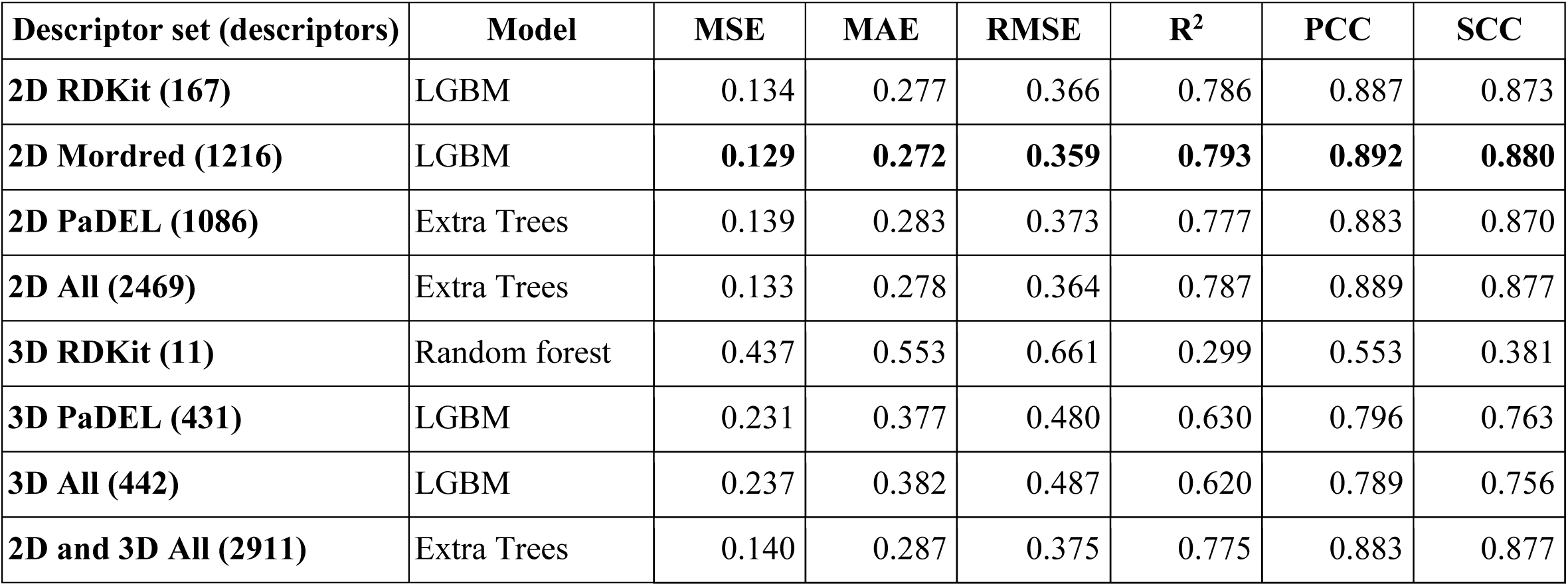
Best-performing regression models for each molecular descriptor set on the Caco2 test dataset. The table reports test set performance across 2D, 3D, and combined descriptor sets using standard regression metrics: MSE, MAE, RMSE, R^2^, PCC, and SCC.

Similar to the PAMPA set, in Caco-2 assay amongst all language models fine-tuned to understand the contextual information from the SMILES, transformer based MoLFormer-XL chemical language model, for the checkpoint MoLFormer-XL-both-10pct, obtained the highest performance (MSE of 0.296, R^2^ of 0.526, and PCC of 0.798) suggesting the effectiveness of the particular language model to fine tune on SMILLES representations. Table S15, list the performance on Caco2 test set for three language models fine-tuned on Caco2 train set. Additionally, models trained on embeddings derived by MoLFormer-XL chemical language model, consistently have superior performance than other two language model-based embeddings (MSE of 0.182, R^2^ of 0.708 and PCC of 0.842), highlighting the superiority of MolFormer language model derived embeddings in predicting the properties of modified peptides with cyclization, including permeability. The test set performances of all the embedding based regressors are listed in Table S16.

To further capitalize on the complementary information embedded across multiple representations, we trained models on combined features of different modalities by selecting top 50 features based on feature importance using random forest. The model with top 50 features achieved the best MSE of 0.184, R^2^ of 0.705 and PCC of 0.841 (Table S17). These models show relatively moderate predictive efficiency when compared to individual modalities like descriptors or fingerprints. Second, we implemented a stacked ensemble model architecture, incorporating atomic features, molecular descriptors, fingerprints and embeddings from language models as input modalities to base learners. The resulting ensemble architecture attained the performance on the test set, with MSE of 0.135, and the highest R^2^ score of 0.783, and PCC of 0.885 (Table 7). Table 8, summarizes the best test set performances by baseline models, language models and stacked ensemble architecture based regressors.

**Table 7.**
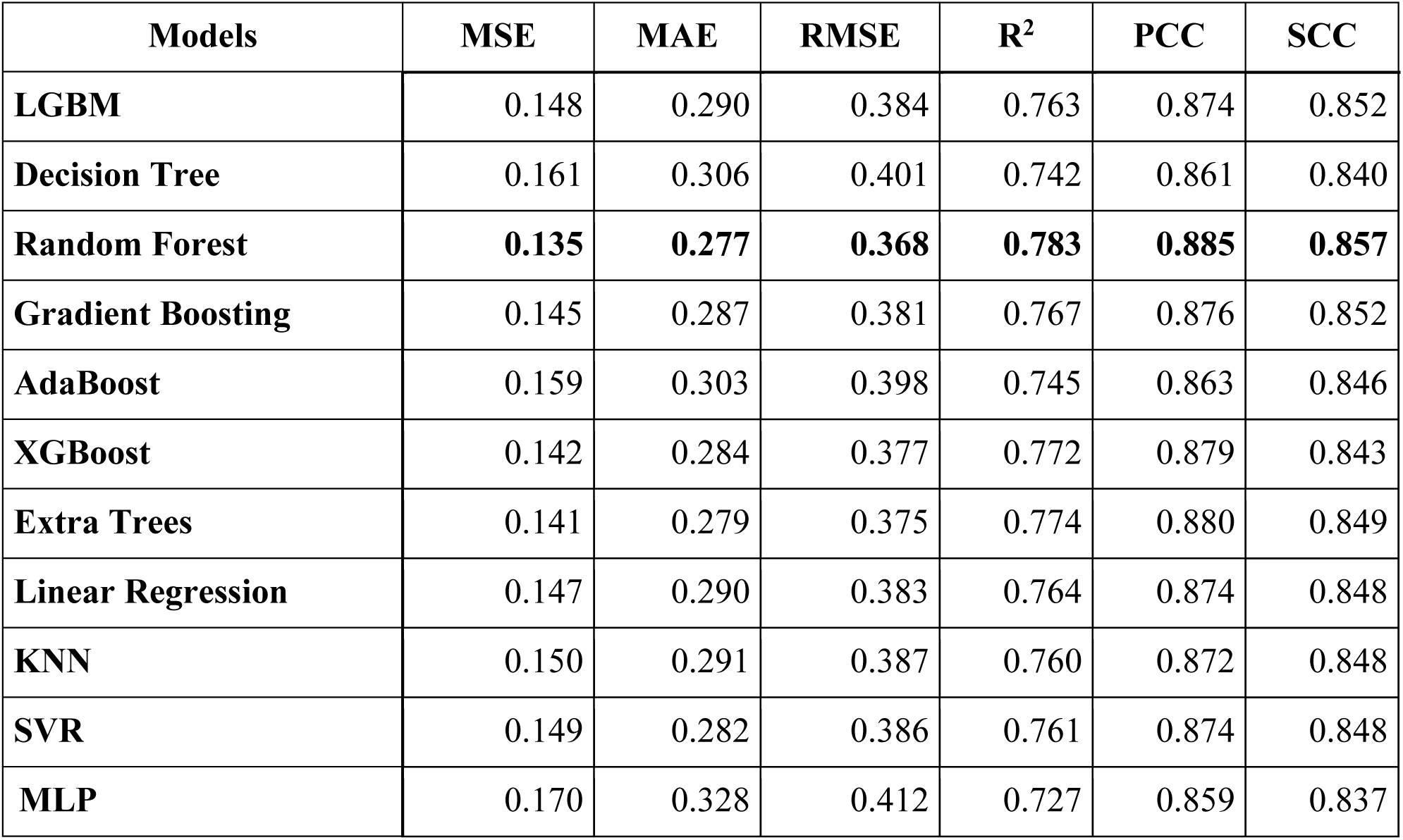
Test set performance metrics for stacked ensemble models for Caco-2 assay.

**Table 8:**
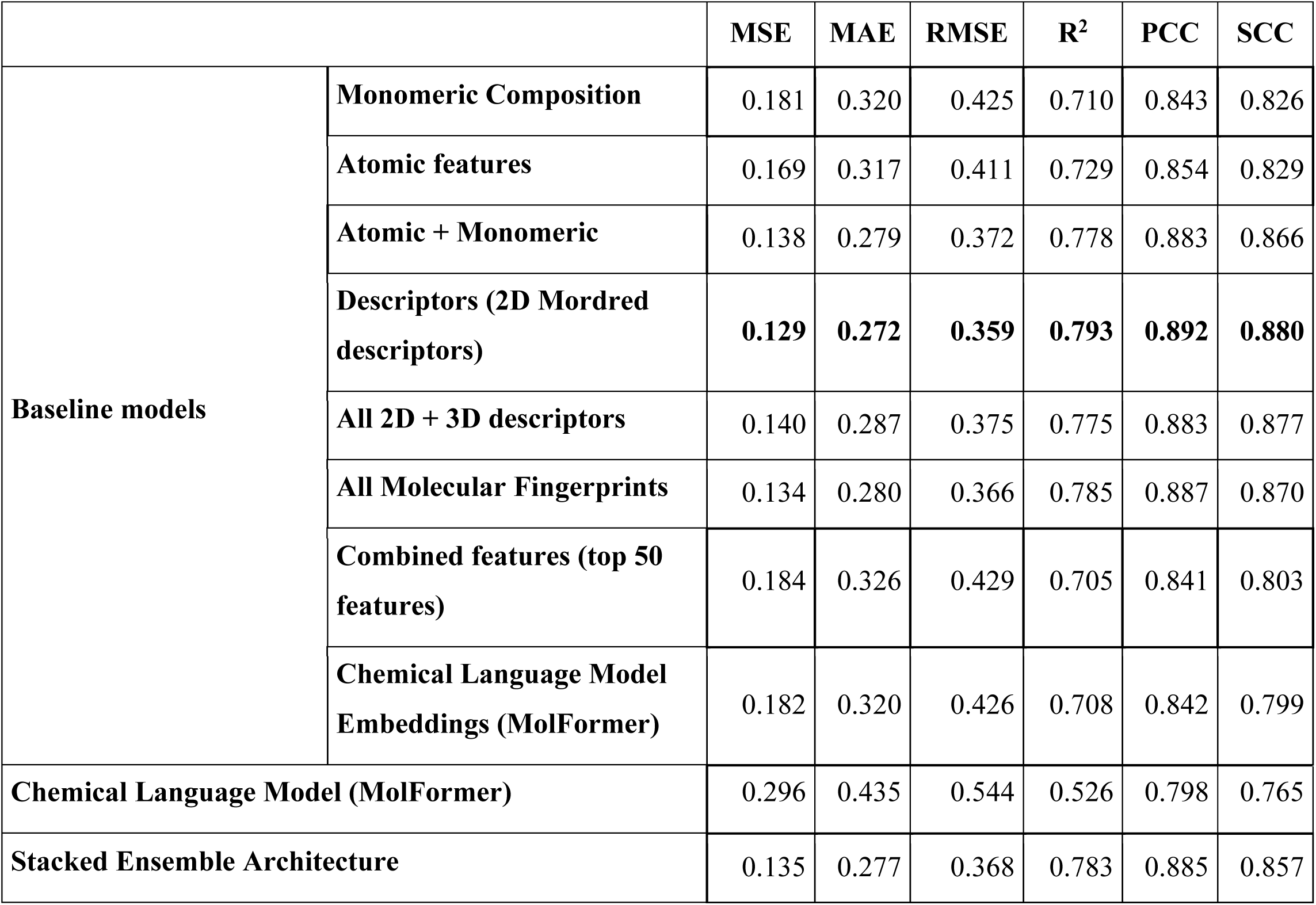
Comparative test set performances of baseline models, chemical language model and ensemble architectures for Caco-2 assay.

To interpret the relative contribution of each representation, an ablation study was performed using the Extra Trees regressor as the final model in stacked architecture (Figure 5B). The exclusion of base models trained on molecular fingerprints resulted in the most significant reduction in R^2^. Additionally, removal of base models trained on cheminformatics derived 2D/3D descriptors also resulted in significantly reduced R^2^ (Figure 5B) and increased MSE, MAE, and RMSE (Figure S9A-C). Embeddings along with atomic features surprisingly show a very slight impact on the modeling efficiency of the stacked architecture.

### Permeability modeling in RRCK assay

The highest performing model to predict permeability derived from RRCK assay with a limited number of datapoints in train set (140 entries) and test set (35 entries) was KlekotaRoth fingerprints, wherein the highest R^2^ of 0.655, PCC score of 0.813 and MSE of 0.160 was achieved. Following that combined 2D descriptors (2367 descriptors after removing the constant features) yielded the R^2^ of 0.645, MSE of 0.164, and PCC of 0.816. The performances of other baseline models, chemical language model and stacked ensemble architecture are all summarised in Table 9. Further, ablation study was conducted for RRCK dataset to evaluate the individual contribution of four molecular feature types in stacked ensemble architecture using AdaBoost as regressor. Excluding models trained on molecular descriptors led to the highest drop in performance (R^2^ score, Figure 5C; MSE, MAE, RMSE, Figure S10A-C), followed by molecular embeddings, indicating their synergistic role in capturing essential molecular information. In contrast, removal of models trained on atomic features or fingerprints show least contrasting influence in the stack architecture.

**Table 9:**
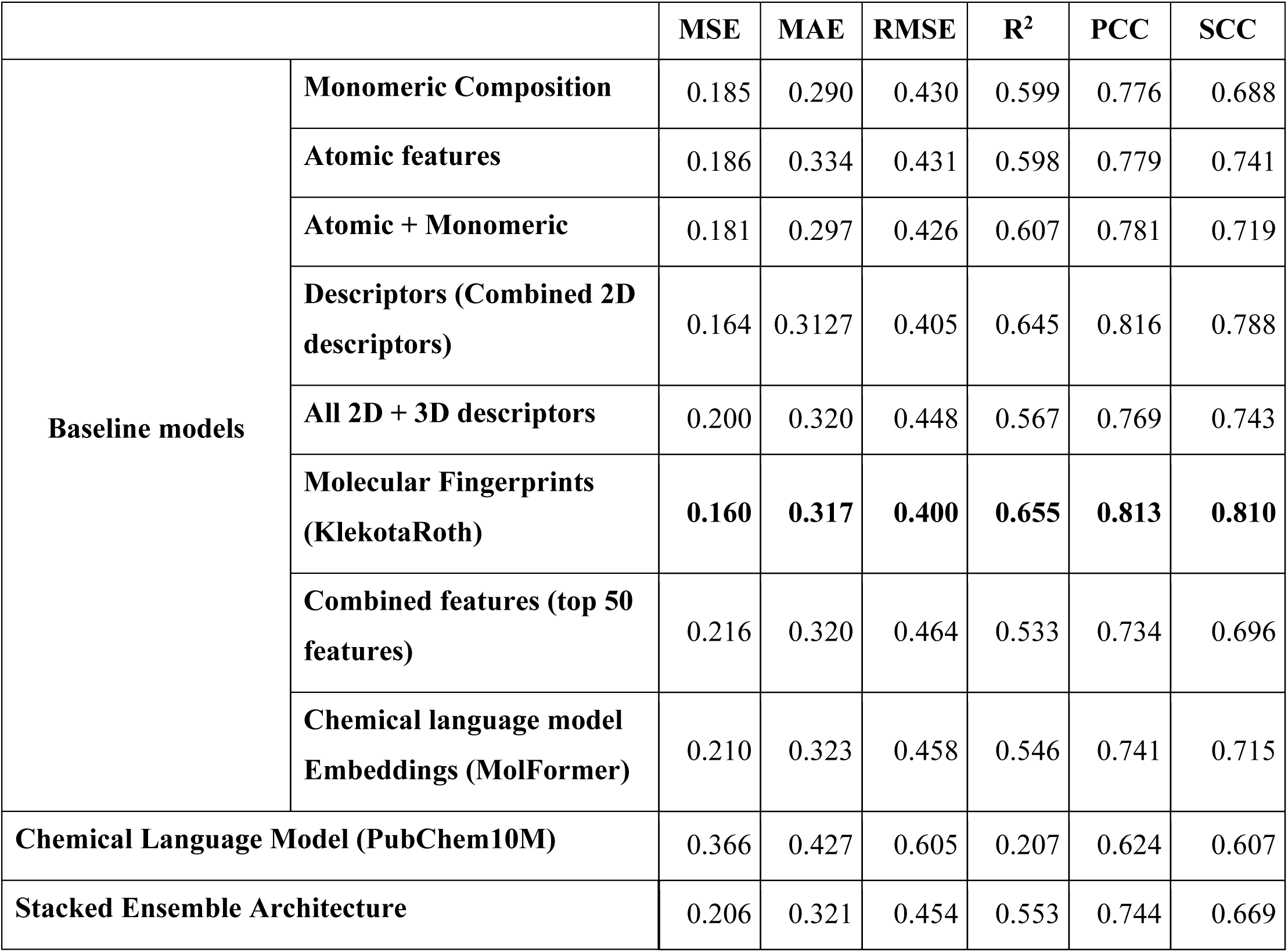
Comparative performances of baseline models, chemical language model and, stacked ensemble architectures for RRCK assay test set.

### Permeability modeling in MDCK assay

Predicting accurate permeability of modified peptides with cyclization in MDCK dataset is limited due to extremely small dataset size upon using the train (51 entries) and test set (13 entries). Stacked ensemble architecture showed the highest performance metric with R^2^ of 0.654, MSE of 0.247, and PCC of 0.894. In baseline models the best performance metric was attained by Morgan fingerprints based regressors (R^2^ of 0.592, MSE of 0.292 and PCC of 0.825). 2D and 3D combined descriptors (3539 descriptors) yielded MSE of 0.350, R^2^ of 0.511 and PCC score of 0.764, whereas, combined atomic and monomeric composition after invariants removal with 44 features yielded a R^2^ of 0.438 and PCC of 0.760. The comparative evaluation of all the different feature based regressors are all shown in Table 10, comparing the performance of a test set by different baseline models and ensemble architecture. Additionally, ablation study conducted for MDCK dataset using XGB regressor as model shows excluding models trained on molecular descriptors led to a prominent reduction in performance (R^2^, Figure 5D; MSE, MAE, RMSE, PCC and SCC Figure S11A-E). Models trained on other features like fingerprints or atomic features have very little effect on overall performance as compared to descriptors. Removal of models trained on embeddings resulted in a rather improved R^2^ score.

**Table 10:**
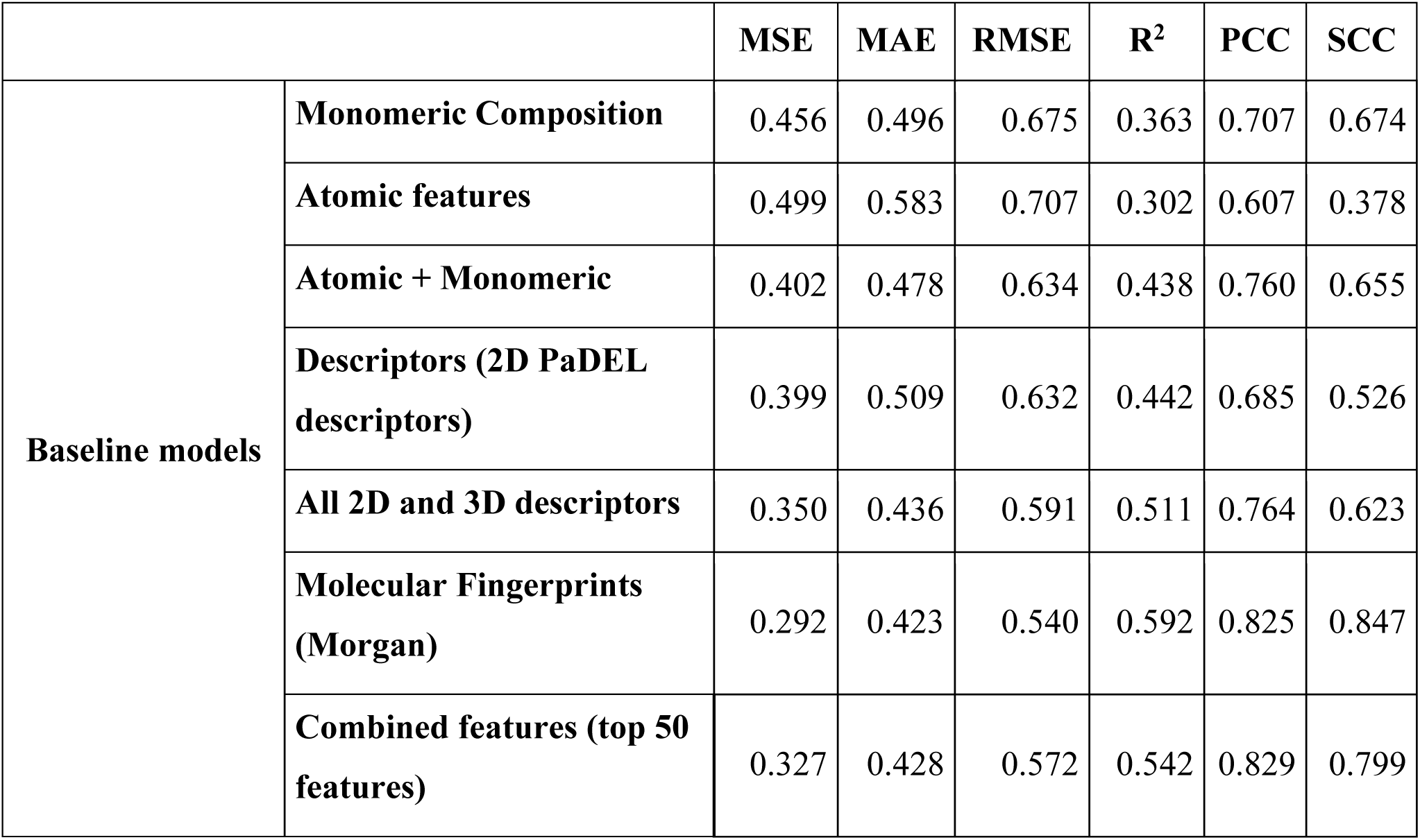
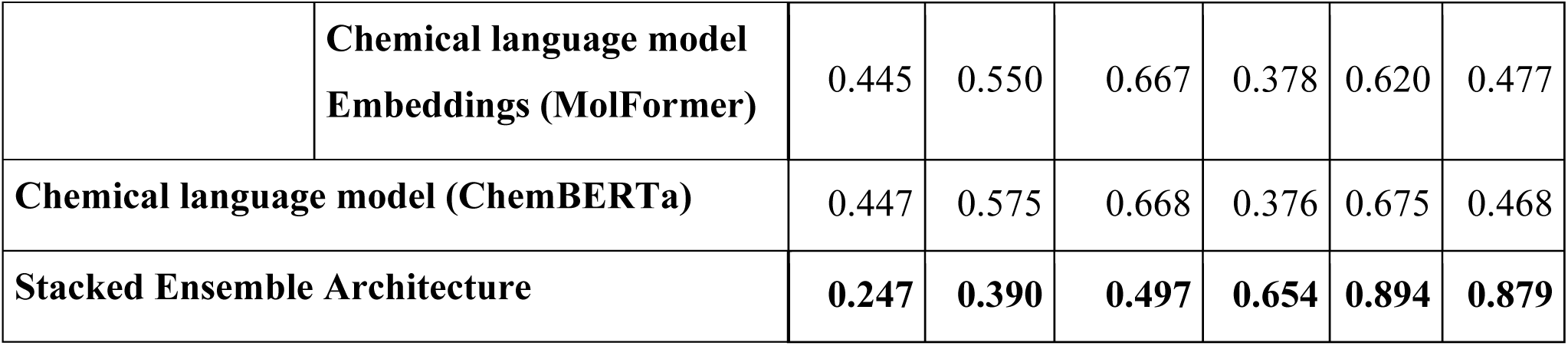
Comparative test performances of baseline models, chemical language model and, stacked ensemble architectures for MDCK assay test set.

### Performance comparison with existing frameworks

Several recent studies including CycPeptMP, Multi_CycGT, PeptideCLM, MuCoCP, CyclePermea, PharmPapp, and CPMP have proposed frameworks to either classify or predict the permeability of cyclic peptides thereby made significant advances in peptide permeability prediction. However, while these methods represent important progress, studies like MuCoCP (R^2^ = 0.338 and MSE = 0.518), CyclePermea (MSE = 0.217, MAE

= 0.334, and PCC = 0.801) proposed generalised prediction frameworks. Additionally, studies like CycPeptMP and PharmPapp use commercial software, or features that are not readily reproducible, thereby restricting their accessibility and wider adoption in academic or resource-constrained environments. Contrary to that, our models (PCPpred), developed exclusively with open-source tools and software, demonstrate superior and/or at least comparable predictive performance across multiple assays (Table 11). Especially, for assays with larger datasets (PAMPA and Caco-2), PCPpred consistently outperformed existing models, achieving higher predictive capability and correlation metrics (PAMPA with R^2^ = 0.685, and PCC = 0.830; Caco-2 with R^2^ = 0.793, and PCC = 0.892) compared to the latest studies CPMP (PAMPA with R^2^ = 0.671, and Caco-2 of R^2^ = 0.746) and PharmPapp (PAMPA with R^2^ = 0.658, Caco-2 with R^2^ = 0.553). Notably, even in case of smaller datasets (RRCK and MDCK), PCPpred maintained competitive performance (RRCK with R^2^ = 0.655, PCC = 0.813; and MDCK with R^2^ = 0.654, PCC = 0.894), thereby closely aligning and/or surpassing previously reported performance metrics. These results collectively highlight the robustness and practical utility of PCPpred as an open-source, reproducible, and high-performing platform for predicting modified peptide permeability across diverse assay types.

**Table 11.**
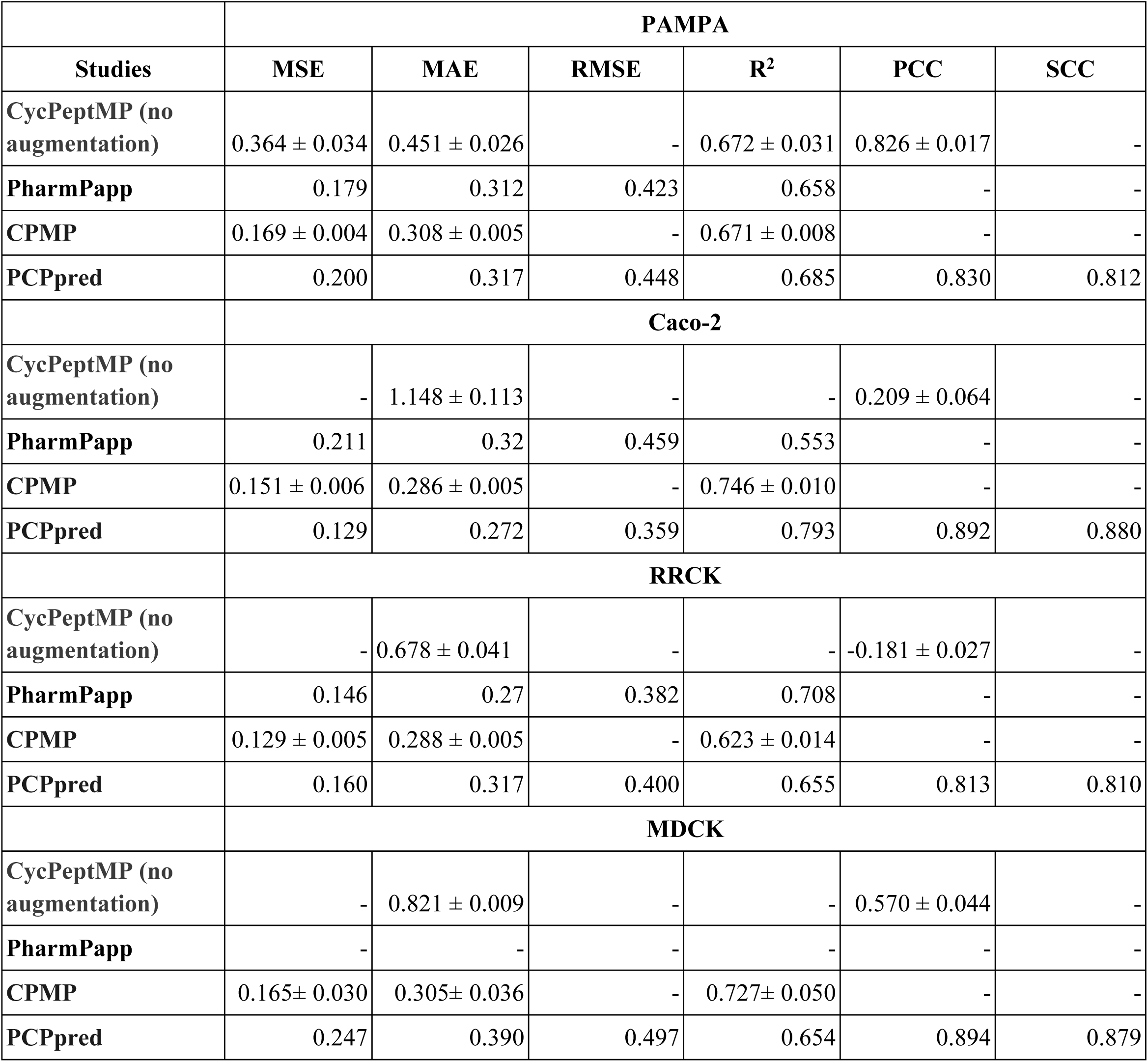
Performance comparison of PCPpred and existing models across permeability assays.

## 4. Discussion

In recent years, a growing shift from small molecules to peptide-based drugs has been witnessed in the therapeutic landscape as evident by the ever-increasing number of FDA approvals being given to peptide-based drugs due to their high specificity, and customizable biological functions [37,38]. This transition has driven the development of various peptide-centric resources such as THPdb, SATPdb, CPPsite, Hemolytik, APD3 and CAMP [39-44]. In parallel, prediction tools such as AntiCP, AntiBP3, AntiFP, AVPpred, AHTpin, and CellPPDmod, have significantly contributed to the in-silico design and functional characterization of the peptides [45-50]. Notably, framework for screening and designing of cell penetrating peptides [51], and discoveries like a novel cell penetrating peptide IMT-P8 for topological delivery of proteins and peptides further highlight the translational potential of peptides in drug delivery [52]. However, the full-scale clinical transition of the peptides remains hindered by challenges like peptide stability, rapid enzymatic degradation and poor half-life in plasma, as reported by tools like PepLife and PlifePred [53,54]. In order to address these challenges, researchers have progressively tried introducing structural modification such as N-methylation, cyclization of peptides or insertion of non-natural residues, to increase the metabolic stability and half-life of the peptides [55,56]. Although advancements are being made in this domain, yet another major challenge that researchers are constantly being faced by in peptide drug development, particularly for oral drug delivery is their limited membrane permeability. Unlike small molecules, modified peptides with cyclization possess high molecular weights, conformational complexity, and structural diversity, thereby making permeability of these peptides difficult to predict [8,57]. Therefore, in this study, we performed a systematic analysis of permeability across multiple assays and integrated diverse molecular features into a predictive machine learning framework, yielding both mechanistic insights and practical advances in peptide design.

The analysis of peptide permeability across multiple assays revealed several consistent and assay-specific patterns that provide valuable insights into the chemical determinants of passive diffusion. Composition analysis showed that canonical residues such as Leu, Ala, Phe, Pro, and Gly were broadly represented across both permeable and non-permeable peptides, suggesting that these residues play a more structural than discriminatory role in determining permeability. By contrast, non-canonical modifications emerged as decisive modulators. N-methylation and bulky hydrophobic substitutions generally enhanced PAMPA permeability, most likely by increasing lipophilicity and shielding hydrogen bond donors, but the same modifications were observed to reduce permeability in Caco-2, reflecting the influence of solubility and efflux mechanisms in cellular contexts. Similarly, polar substitutions such as Asp-piperidide and Ser(tBu) showed improved permeability in Caco-2, underscoring that cell-based transport may sometimes favour polar chemical features. Descriptor-level patterns reinforced the findings by demonstrating that permeability depends on achieving a balance between moderate lipophilicity and controlled polarity, often realized through intramolecular hydrogen bonding or shielding, rather than extreme hydrophobicity.

Furthermore, descriptor correlation analysis revealed that while Caco-2, RRCK, and MDCK assays were highly concordant with one another, PAMPA exhibited distinct and, in some cases, inverse descriptor permeability relationships. This divergence indicates that permeability mechanisms captured in PAMPA may differ fundamentally from those in cellular assays. As a result, models trained exclusively on PAMPA data might not fully capture the trends observed in Caco-2 or MDCK, and thus may not serve as direct surrogates for predicting cellular permeability. Consistent with this, pooled predictive models trained across all assays achieved only moderate accuracy, while performance improved substantially when assay identity was explicitly included as a meta-feature. These results emphasize that permeability prediction is assay-contextual, and highlight the necessity of developing individual models tailored to each assay rather than relying solely on generalized predictors.

Building on these insights, we systematically evaluated a wide range of molecular representations for predictive modeling. Individual feature classes showed varying contributions across assays like, atomic features provided consistent but modest predictive power, fingerprints such as Klekota-Roth were particularly effective in small datasets like RRCK, and chemical language model embeddings (MolFormer-XL) emerged as highly predictive in PAMPA. Classical physicochemical descriptors demonstrated strong predictive ability in Caco-2, where substructural and polarity-related properties were closely aligned with permeability trends. Importantly, ensemble models that integrated these heterogeneous features consistently achieved superior performance in large datasets, with PAMPA and Caco-2 showing marked comparable performance and gains over single-modality baselines. However, the benefits of ensemble architectures were diminished in extremely small datasets such as RRCK, where simpler modalities like fingerprint-based models sometimes outperformed stacked models, underscoring the importance of data volume and diversity in determining the success of ensemble learning.

Taken together, these findings confirm that the proposed machine learning framework (stacked ensemble framework) provides accurate and interpretable predictions of modified peptide permeability especially in larger datasets by leveraging complementary strengths of atomic descriptors, classical physicochemical features, fingerprints, and transformer embeddings. The models achieved comparable and/or outperform performances achieved by latest prediction studies of peptides such as CPMP and thereby establish a scalable approach capable of adapting across assays of varying sizes and complexity. To ensure these advances are broadly accessible, we implemented a user-friendly web server that enables researchers to design peptides using MAP (modification and annotations of proteins) notation, which can then directly be used to predict permeability or can be converted in widely used SMILES notation. Input of MAP format is automatically converted to SMILES sequence, followed by generating molecular features, and obtaining assay-specific permeability predictions with visual summaries. By providing a streamlined platform that eliminates the need for specialized software or coding expertise, the server empowers chemists and biologists to incorporate permeability predictions directly into their design workflows during pre-testing phase. This integration of mechanistic insights, predictive modeling, and accessible implementation offers a comprehensive framework for accelerating peptide optimization and drug discovery.

## 5. Conclusion

In summary, this study presents a robust and interpretable analysis of peptides and their permeability obtained through different assays, and machine learning frameworks tailored for predicting the passive permeability of modified peptides with cyclization, across multiple biological assays. By integrating heterogeneous molecular representations within an ensemble architecture, the approach effectively captures the multifactorial nature of peptide physicochemical properties like permeability. Its consistent superiority over baseline models and existing methods, particularly in large datasets like PAMPA and Caco-2, highlights its scalability and predictive strength. Furthermore, the development of an open-access web server featuring a human-readable MAP design interface and automated prediction pipeline ensures broad usability, enabling researchers to evaluate novel peptides without reliance on proprietary software. This work not only advances the field of peptide pharmacokinetics modeling but also lays a practical foundation for accessible, data-driven peptide design and optimization.

## Supporting information

Supplementary tables from Table S1 to Table S17, supplementary figures from Figure S1 to Figure S11, all in order as they appear in the manuscript.

## 7. Data Availability Statement

All the data and results used and/or generated in this study are available at the “PCPpred” webserver, https://webs.iiitd.edu.in/raghava/pcppred; and GitHub repository, https://github.com/raghavagps/pcppred.

## 9. Supporting Information

Supplementary tables from Table S1 to Table S17, supplementary figures from Figure S1 to Figure S11, all in order as they appear in the manuscript (Doc).

## 10. Corresponding Author Information

Prof. Gajendra P. S. Raghava: Phone: 011-26907444

## 11. Author Contributions

AS and GPSR conceptualized the idea. AS and GPSR collected and processed the datasets. AS developed and implemented the prediction models/framework. AS and GPSR analysed the results. AS and PSG created the front-end of the user interface. AS wrote the scripts and back-end of the web server. AS, and GPSR penned the manuscript. AS, GPSR and PSG edited the manuscript. GPSR conceived and coordinated the project. The authors have read and approved the final manuscript.

## 12. Acknowledgements

The authors gratefully acknowledge the financial support provided by the Department of Biotechnology (DBT). The authors are also thankful to the University Grants Commission (UGC), Council of Scientific and Industrial Research (CSIR) for the fellowship. They also extend their sincere thanks to the Department of Computational Biology at the Indraprastha Institute of Information Technology Delhi (IIIT-Delhi) for providing the necessary computational infrastructure and research facilities. Furthermore, they would like to acknowledge Bioicons.com, draw.io and NIH BIOART for the creation of the figures utilized in this work.

## 13. Abbreviations

PAMPA: Parallel Artificial Membrane Permeability Assay
RRCK: Ralph Russ Canine Kidney
MDCK: Madin-Darby Canine Kidney
KDE: Kernel density estimation
LGBM: Light Gradient Boosting Machine
XGBoost: eXtreme Gradient Boosting
AdaBoost: Adaptive Boosting
SVR: Support Vector Regressor
KNN: K-Neighbors Regressor
MLP: Multi-Layer Perceptron
MSE: Mean Squared Error
RMSE: Root Mean Squared Error
MAE: Mean Absolute Error
R^2^: Coefficient of Determination
PCC: Pearson Correlation Coefficient
SCC: Spearman Correlation Coefficient

## [15] For Table of Contents Only

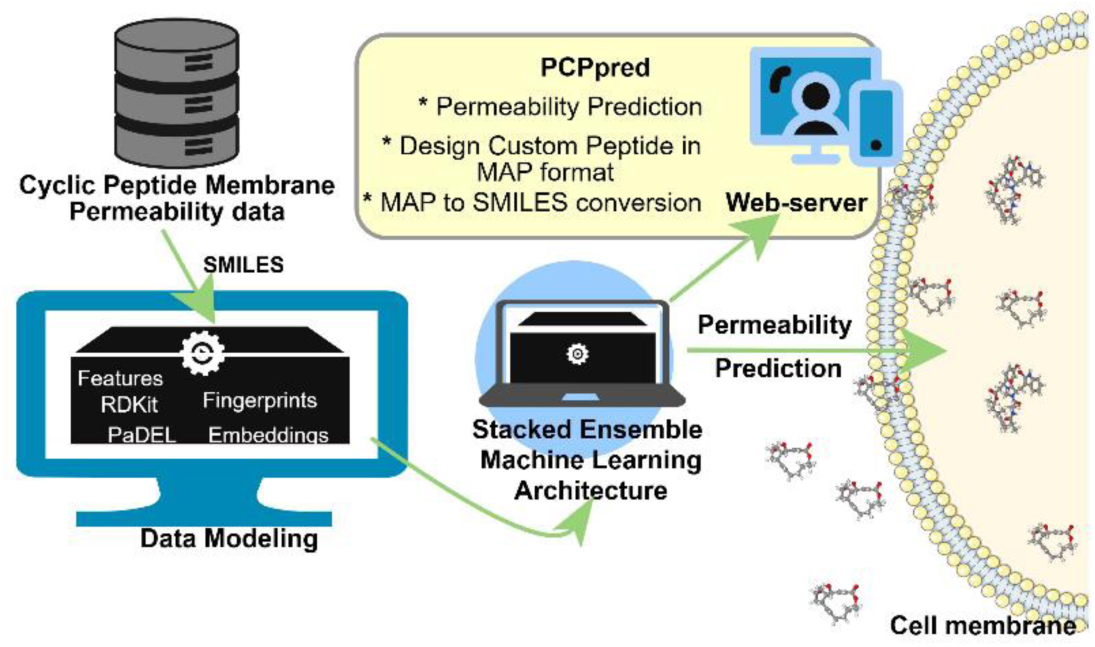

