## Supplementary tables from Table S1 to Table S17, supplementary figures from Figure S1 to Figure S11, all in order as they appear in the manuscript. for "PCPpred: Prediction of Chemically Modified Peptide Permeability Across Multiple Assays for Oral Delivery"

**Mailing Address of Authors**

Akshay Shendre:

ORCID ID: <https://orcid.org/0009-0006-1239-5881>

Pushpendra Singh Gahlot:

ORCID ID: <https://orcid.org/0009-0004-9868-358X>

Gajendra P. S. Raghava (GPSR):

ORCID ID: <https://orcid.org/0000-0002-8902-2876>

***Corresponding Author**

Prof. Gajendra P. S. Raghava

Head and Professor, Department of Computational Biology

Indraprastha Institute of Information Technology, Delhi

Okhla Phase 3, New Delhi, India - 110020

Website: <http://webs.iiitd.edu.in/raghava/>

**Supplementary Tables**

**Table S1:** Inter-assay pairwise correlation coefficient (r) matrix of descriptor-permeability relationships across Caco-2, PAMPA, RRCK, and MDCK datasets.

| **Assay** | **Caco-2** | **PAMPA** | **RRCK** | **MDCK** |
| --- | --- | --- | --- | --- |
| **Caco-2** | 1.00 | -0.77 | 0.96 | 0.97 |
| **PAMPA** | -0.77 | 1.00 | -0.81 | -0.72 |
| **RRCK** | 0.96 | -0.81 | 1.00 | 0.98 |
| **MDCK** | 0.97 | -0.72 | 0.98 | 1.00 |

**Table S2:** Permeability variations of sample peptides shared across multiple assays (PAMPA, Caco-2, RRCK, and MDCK).

| **ID** | **PAMPA** | **Caco2** | **RRCK** | **MDCK** |
| --- | --- | --- | --- | --- |
| **S1** | -5.5 | -4.9 | -5.2 | -5.0 |
| **S2** | -6.4 | -5.5 | -5.8 | -5.7 |
| **S3** | -5.7 | -5.1 | -5.9 | - |
| **S4** | -5.3 | -5.9 | -6.2 | - |
| **S5** | -5.6 | -6.6 | -6.4 | - |
| **S6** | -5.8 | -6.6 | -6.0 | - |
| **S7** | -5.8 | -6.8 | -6.1 | - |
| **S8** | -6.9 | -7.2 | - | - |
| **S9** | -5.1 | -5.6 | - | - |
| **S10** | -6.1 | - | -6.1 | - |
| **S11** | -6.3 | - | -6.8 | - |
| **S12** | -7.8 | - | - | -6.4 |
| **S13** | -4.6 | - | - | -5.2 |

**Table S3:** SMILES representations of peptides corresponding to IDs in Table S2.

| ID | SMILES |
| --- | --- |
| **S1** | CC(C)C[C@@H]1NC(=O)[C@H](Cc2ccc(O)cc2)N(C)C(=O)[C@H]2CCCN2C(=O)[C@H](CC(C)C)NC(=O)[C@H](CC(C)C)N(C)C(=O)[C@@H](CC(C)C)N(C)C1=O |
| **S2** | CC(C)C[C@@H]1NC(=O)[C@H](Cc2ccc(O)cc2)NC(=O)[C@H]2CCCN2C(=O)[C@H](CC(C)C)NC(=O)[C@H](CC(C)C)NC(=O)[C@@H](CC(C)C)NC1=O |
| **S3** | CC[C@H](C)[C@@H]1NC(=O)[C@@H]2CCCN2C(=O)[C@H](Cc2ccccc2)NC(=O)[C@H](C(C)(C)C)NC(=O)c2csc(n2)[C@H]([C@H](C)CC)NC(=O)[C@@H]2CCCN2C1=O |
| **S4** | CC[C@H](C)[C@@H]1NC(=O)[C@@H]2CCCN2C(=O)[C@H](Cc2ccccc2)NC(=O)[C@H](C)NC(=O)c2csc(n2)[C@H]([C@H](C)CC)NC(=O)[C@@H]2CCCN2C1=O |
| **S5** | CC(C)C[C@@H]1NC(=O)[C@@H](CC(C)C)NC(=O)[C@@H](CC(C)C)NC(=O)[C@@H](CC(C)C)NC(=O)[C@H]2CCCN2C(=O)[C@H](Cc2ccc(O)cc2)NC1=O |
| **S6** | CC(C)C[C@@H]1NC(=O)[C@H](CC(C)C)NC(=O)[C@@H](CC(C)C)NC(=O)[C@@H](CC(C)C)NC(=O)[C@H]2CCCN2C(=O)[C@H](Cc2ccc(O)cc2)NC1=O |
| **S7** | CC(C)C[C@@H]1NC(=O)[C@@H](CC(C)C)NC(=O)[C@H](CC(C)C)NC(=O)[C@@H](CC(C)C)NC(=O)[C@H]2CCCN2C(=O)[C@H](Cc2ccc(O)cc2)NC1=O |
| **S8** | CC[C@H](C)[C@@H]1NC(=O)[C@H](Cc2ccccc2)N(C)C(=O)[C@H]([C@@H](C)O)NC(=O)[C@@H](C)NC(=O)C[C@@H](C(=O)N(C)[C@@H](Cc2ccccc2)C(=O)N(C)[C@@  H](Cc2ccccc2)C(=O)N[C@@H](C)C(=O)N2CCCCC2)NC(=O)[C@H](CC(C)C)N(C)C(=O)[C@H](CC(C)C)N(C)C(=O)[C@H](Cc2ccccc2)N(C)C(=O)[C@H](CC(C)C)N(C)C(=O)[C  @H](C)N(C)C1=O |
| **S9** | CC[C@H](C)[C@@H]1NC(=O)[C@H](CC(C)C)N(C)C(=O)[C@H](C)N(C)C(=O)[C@H]([C@@H](C)O)NC(=O)[C@H](Cc2ccccc2)N(C)C(=O)[C@H](C)N(C)C(=O)[C@@H](C)N  (C)C(=O)C[C@@H](C(=O)N(C)[C@@H](Cc2ccccc2)C(=O)N(C)[C@@H](Cc2ccccc2)C(=O)N[C@@H](C)C(=O)N2CCCCC2)NC(=O)[C@H](C(C)C)N(C)C(=O)[C@H](CC(C)C)  N(C)C1=O |
| **S10** | C/C=C/C[C@@H](C)[C@@H](O)[C@H]1C(=O)N[C@@H](C(C)C)C(=O)N(C)CC(=O)N(C)[C@@H](CC(C)C)C(=O)N[C@@H](C(C)C)C(=O)N(C)[C@@H](CC(C)C)C(=O)N[C  @@H](C)C(=O)N[C@H](C)C(=O)N(C)[C@@H](CC(C)C)C(=O)N(C)[C@@H](CC(C)C)C(=O)N(C)[C@@H](C(C)C)C(=O)N1C |
| **S11** | C/C=C/C[C@@H](C)[C@H]1OC(=O)[C@H](C(C)C)N(C)C(=O)[C@H](CC(C)C)N(C)C(=O)[C@H](CC(C)C)N(C)C(=O)[C@@H](C)NC(=O)[C@H](C)NC(=O)[C@H](CC(C)C)N  (C)C(=O)[C@H](C(C)C)NC(=O)[C@H](CC(C)C)N(C)C(=O)CN(C)C(=O)[C@H](CC)NC(=O)[C@H]1NC |
| **S12** | CC(=O)N1CCC[C@@H]1C(=O)N(C)[C@H](CC(C)C)C(=O)N[C@@H]1C(=O)N[C@H](CC(C)C)C(=O)N[C@H](CC(C)C)C(=O)N[C@@H](CC(C)C)C(=O)N[C@@H](C)C(=O)N  [C@@H](CC(C)C)C(=O)N2CCC[C@@H]2C(=O)O[C@@H]1C |
| **S13** | CC(C)C[C@@H]1NC(=O)[C@H](C)N(C)C(=O)[C@H]2CCCN2C(=O)[C@H](CC(C)C)NC(=O)[C@H](C)N(C)C(=O)[C@H](CC(C)C)NC(=O)[C@H](C)N(C)C(=O)[C@H]2CCCN2  C(=O)[C@H](CC(C)C)NC(=O)[C@H](C)N(C)C1=O |

**Table S4.** Statistical comparison of physicochemical descriptors between permeable and non-permeable peptides in the PAMPA train and test datasets.

| **Descriptors** | **PAMPA Train dataset** | | **PAMPA Test dataset** | |
| --- | --- | --- | --- | --- |
|  | **t-statistic** | **p-value** | **t-statistic** | **p-value** |
| **PEOE_VSA8** | -16.29252276 | 8.46E-58 | -7.237538504 | 9.17E-13 |
| **AATSC.6** | 15.14615735 | 2.94E-50 | 7.468573264 | 1.87E-13 |
| **minsOH** | -11.28095992 | 3.93E-29 | -6.518822366 | 1.03E-10 |
| **CrippenLogP** | -10.05742501 | 2.33E-23 | -4.633607813 | 4.32E-06 |
| **MolecularId.7** | 9.894678612 | 1.11E-22 | 4.973325689 | 8.33E-07 |
| **MolLogP** | -9.784582366 | 3.24E-22 | -4.906370399 | 1.16E-06 |
| **PEOE_VSA6** | -8.805545007 | 2.22E-18 | -4.170010468 | 3.40E-05 |
| **BCUT2D_MWLOW** | 7.94946701 | 2.70E-15 | 3.795870462 | 0.000159949859 |
| **BCUT.7** | -7.287519101 | 4.35E-13 | -3.265668548 | 0.001155169582 |
| **RDF20m** | -7.109584014 | 1.40E-12 | -3.297309956 | 0.001014023355 |
| **TopologicalCharge.12** | -5.672242839 | 1.55E-08 | -3.258099297 | 0.00117233131 |
| **Chi.9** | 5.237634932 | 1.73E-07 | 2.987415757 | 0.00289915363 |
| **minssCH2** | 4.190623836 | 2.85E-05 | 2.805067535 | 0.005133709009 |
| **RotatableBondsRatio** | 3.884817818 | 0.0001047019106 | 2.561982911 | 0.01061092324 |
| **EtaVEMCount.3** | 3.635851522 | 0.0002818191018 | 1.573595386 | 0.1159762525 |
| **E1p** | -3.114813191 | 0.001857976311 | -1.84427747 | 0.06554236045 |
| **RPCS** | 2.937905422 | 0.003326184568 | 1.915492389 | 0.05590877939 |
| **XLogP** | 2.474044965 | 0.01342071803 | 2.203100595 | 0.02791226296 |
| **TDB2i** | -1.476773292 | 0.1398938121 | -1.304259718 | 0.1928254708 |
| **TDB1i** | -0.3277037519 | 0.7431700411 | -0.8089718962 | 0.418968433 |

**Table S5.** Statistical comparison of physicochemical descriptors between permeable and non-permeable peptides in the Caco2 train and test datasets.

| **Descriptors** | **Caco2 Train dataset** | | **Caco2 Test dataset** | |
| --- | --- | --- | --- | --- |
|  | **t-statistic** | **p-value** | **t-statistic** | **p-value** |
| **AtomTypeEState.252** | -10.25122958 | 3.15E-23 | -4.172614965 | 4.49E-05 |
| **TDB9s** | 10.05294835 | 2.63E-22 | 4.627624631 | 7.60E-06 |
| **minwHBa** | -9.906374099 | 5.99E-22 | -3.104881585 | 0.002202718668 |
| **TDB10s** | 9.556628556 | 1.68E-20 | 4.994489802 | 1.47E-06 |
| **maxHBint10** | 9.385032618 | 4.38E-20 | 4.515526722 | 1.02E-05 |
| **AtomTypeEState.94** | -8.439693947 | 1.51E-16 | -3.035737478 | 0.002715394795 |
| **minssCH2** | -8.455023852 | 1.53E-16 | -3.679403545 | 0.0003090776405 |
| **TDB8s** | 8.319054302 | 4.21E-16 | 4.032834664 | 8.49E-05 |
| **DPSA-3** | 7.736236636 | 2.97E-14 | 3.354189445 | 0.0009480956898 |
| **fr_Ndealkylation2** | 7.488165623 | 1.68E-13 | 2.445889817 | 0.01521796057 |
| **AATS8e** | 7.499780707 | 1.92E-13 | 4.003320946 | 9.43E-05 |
| **FNSA-2** | -7.192814803 | 1.41E-12 | -3.129521894 | 0.001997271983 |
| **PNSA-1** | 6.980191874 | 5.76E-12 | 3.235468464 | 0.001395667434 |
| **maxHBint6** | 6.749592159 | 2.53E-11 | 4.026188345 | 7.57E-05 |
| **nHBint10** | 6.552650811 | 9.23E-11 | 2.708295415 | 0.007236892963 |
| **maxHssNH** | 6.380378697 | 2.75E-10 | 3.539629824 | 0.0004793900007 |
| **minsCH3** | -5.833014283 | 7.87E-09 | -2.079987742 | 0.03857434114 |
| **MolLogP** | -5.162553102 | 2.94E-07 | -2.556155856 | 0.01117842954 |
| **SLogP** | -5.162553102 | 2.94E-07 | -2.556155856 | 0.01117842954 |
| **CrippenLogP** | -4.784229658 | 1.98E-06 | -2.385046977 | 0.01782752604 |

**Table S6.** Predictive performance of machine learning models on the PAMPA test dataset using atomic features alone versus a combination of atomic features and monomeric composition. Evaluation metrics include MSE, MAE, RMSE, R², PCC, and SCC.

|  | **Atomic features** | | | | | | **Atomic features + monomeric composition** | | | | | |
| --- | --- | --- | --- | --- | --- | --- | --- | --- | --- | --- | --- | --- |
| **Models** | **MSE** | **MAE** | **RMSE** | **R^2^** | **PCC** | **SCC** | **MSE** | **MAE** | **RMSE** | **R^2^** | **PCC** | **SCC** |
| **LGBM** | 0.309 | 0.406 | 0.556 | 0.514 | 0.718 | 0.701 | 0.254 | 0.367 | 0.504 | 0.601 | 0.776 | 0.750 |
| **Decision Trees** | 0.309 | 0.391 | 0.556 | 0.514 | 0.722 | 0.716 | 0.299 | 0.389 | 0.547 | 0.530 | 0.739 | 0.698 |
| **Random Forest** | 0.289 | 0.387 | 0.537 | 0.546 | 0.740 | 0.720 | 0.258 | 0.364 | 0.508 | 0.594 | 0.771 | 0.738 |
| **Gradient Boosting** | 0.341 | 0.436 | 0.584 | 0.464 | 0.686 | 0.664 | 0.290 | 0.405 | 0.539 | 0.544 | 0.747 | 0.714 |
| **AdaBoost** | 0.536 | 0.585 | 0.732 | 0.157 | 0.483 | 0.443 | 0.478 | 0.553 | 0.691 | 0.248 | 0.563 | 0.519 |
| **XGBoost** | **0.283** | **0.383** | **0.532** | **0.554** | **0.746** | **0.724** | **0.226** | **0.342** | **0.475** | **0.645** | **0.803** | **0.775** |
| **Extra Trees** | 0.291 | 0.386 | 0.539 | 0.543 | 0.739 | 0.718 | 0.261 | 0.368 | 0.511 | 0.589 | 0.771 | 0.729 |
| **Linear Regression** | 0.496 | 0.528 | 0.704 | 0.220 | 0.470 | 0.487 | 0.375 | 0.440 | 0.612 | 0.411 | 0.651 | 0.681 |
| **KNN** | 0.335 | 0.411 | 0.579 | 0.473 | 0.698 | 0.681 | 0.334 | 0.422 | 0.578 | 0.475 | 0.695 | 0.664 |
| **SVR** | 0.381 | 0.433 | 0.617 | 0.401 | 0.642 | 0.638 | 0.302 | 0.391 | 0.549 | 0.526 | 0.729 | 0.708 |
| **MLP** | 0.346 | 0.425 | 0.588 | 0.456 | 0.677 | 0.661 | 0.316 | 0.398 | 0.562 | 0.503 | 0.725 | 0.727 |

**Table S7.** Performance comparison of machine learning regressors on the PAMPA test dataset using monomer composition and their natural amino acid analogs as input features. Test set performance metrics include MSE, MAE, RMSE, R², PCC, and SCC to evaluate permeability prediction.

|  | **Monomer Composition** | | | | | | **Natural analog monomeric Composition** | | | | | |
| --- | --- | --- | --- | --- | --- | --- | --- | --- | --- | --- | --- | --- |
| **Models** | **MSE** | **MAE** | **RMSE** | **R^2^** | **PCC** | **SCC** | **MSE** | **MAE** | **RMSE** | **R^2^** | **PCC** | **SCC** |
| **Extra Trees** | 0.292 | 0.388 | 0.541 | 0.540 | 0.741 | 0.711 | 0.407 | 0.469 | 0.638 | 0.360 | 0.602 | 0.544 |
| **LGBM** | 0.308 | 0.407 | 0.555 | 0.516 | 0.720 | 0.701 | 0.407 | 0.472 | 0.638 | 0.360 | 0.600 | 0.545 |
| **XGBoost** | **0.258** | **0.372** | **0.508** | **0.595** | **0.772** | **0.741** | **0.403** | **0.465** | **0.634** | **0.367** | **0.607** | **0.546** |
| **Decision Tree** | 0.327 | 0.405 | 0.572 | 0.486 | 0.713 | 0.685 | 0.413 | 0.471 | 0.643 | 0.350 | 0.595 | 0.535 |
| **Random Forest** | 0.281 | 0.383 | 0.530 | 0.558 | 0.748 | 0.717 | 0.406 | 0.469 | 0.637 | 0.362 | 0.602 | 0.545 |
| **Gradient Boosting** | 0.340 | 0.436 | 0.583 | 0.466 | 0.695 | 0.670 | 0.431 | 0.494 | 0.656 | 0.323 | 0.569 | 0.515 |
| **AdaBoost** | 0.492 | 0.565 | 0.701 | 0.227 | 0.550 | 0.499 | 0.535 | 0.587 | 0.732 | 0.158 | 0.451 | 0.432 |
| **SVR** | 0.318 | 0.398 | 0.564 | 0.500 | 0.710 | 0.699 | 0.438 | 0.472 | 0.662 | 0.311 | 0.570 | 0.524 |
| **Linear Regression** | 0.368 | 0.443 | 0.607 | 0.422 | 0.652 | 0.666 | 0.493 | 0.533 | 0.703 | 0.224 | 0.474 | 0.442 |
| **KNN** | 0.346 | 0.428 | 0.588 | 0.456 | 0.683 | 0.652 | 0.546 | 0.548 | 0.739 | 0.142 | 0.510 | 0.460 |
| **MLP** | 0.374 | 0.423 | 0.611 | 0.412 | 0.673 | 0.683 | 0.433 | 0.486 | 0.658 | 0.318 | 0.567 | 0.526 |

**Table S8.** Predictive performances of best-performing regressors trained on different molecular fingerprint types for the PAMPA test set. Test set performance metrics reported include MSE, MAE, RMSE, R², PCC, and SCC.

| **Fingerprint Type** | **Model** | **MSE** | **MAE** | **RMSE** | **R^2^** | **PCC** | **SCC** |
| --- | --- | --- | --- | --- | --- | --- | --- |
| **All fingerprints** | XGBoost | **0.212** | **0.332** | **0.461** | **0.666** | **0.816** | **0.799** |
| **AtomPairs2D** | Random Forest | 0.466 | 0.521 | 0.683 | 0.267 | 0.517 | 0.463 |
| **Count AtomPairs2D** | Extra Trees | 0.233 | 0.343 | 0.482 | 0.634 | 0.797 | 0.785 |
| **Count KlekotaRoth** | Extra Trees | 0.236 | 0.349 | 0.486 | 0.628 | 0.793 | 0.778 |
| **Count Morgan** | XGBoost | **0.226** | **0.342** | **0.475** | **0.645** | **0.803** | **0.787** |
| **Count Substructure** | XGBoost | 0.251 | 0.353 | 0.501 | 0.605 | 0.778 | 0.766 |
| **EState** | Gradient Boosting | 0.487 | 0.541 | 0.698 | 0.235 | 0.485 | 0.397 |
| **Extended** | XGBoost | 0.365 | 0.452 | 0.604 | 0.426 | 0.654 | 0.594 |
| **Fingerprinter** | XGBoost | 0.376 | 0.464 | 0.613 | 0.408 | 0.640 | 0.558 |
| **GraphOnly** | Random Forest | 0.417 | 0.485 | 0.646 | 0.344 | 0.588 | 0.516 |
| **KlekotaRoth** | XGBoost | 0.292 | 0.398 | 0.540 | 0.542 | 0.736 | 0.709 |
| **MACCS** | Random Forest | 0.344 | 0.430 | 0.587 | 0.459 | 0.678 | 0.657 |
| **Morgan** | XGBoost | 0.252 | 0.362 | 0.502 | 0.603 | 0.777 | 0.765 |
| **PubChem** | XGBoost | 0.368 | 0.456 | 0.606 | 0.422 | 0.650 | 0.577 |
| **Substructure** | Random Forest | 0.473 | 0.519 | 0.688 | 0.256 | 0.509 | 0.431 |

**Table S9.** Comparative test set performance of different pre-trained molecular language models, MoLFormer XL both-10pct, PubChem10M SMILES BPE 450K, and ChemBERTa-zinc-base-v1, fine-tuned on the PAMPA training set. Evaluation on the test set was conducted using standard regression metrics: MSE, MAE, RMSE, R², PCC, and SCC.

| **Metrics** | **MolFormer_XL_both-10pct** | **PubChem10M_SMILES_BPE_450K** | **ChemBERTa-zinc-base-v1** |
| --- | --- | --- | --- |
| **MSE** | 0.336 | **0.281** | 0.298 |
| **MAE** | 0.465 | **0.376** | 0.385 |
| **RMSE** | 0.580 | **0.530** | 0.546 |
| **R^2^** | 0.471 | **0.558** | 0.532 |
| **PCC** | **0.801** | 0.762 | 0.752 |
| **SCC** | **0.787** | 0.744 | 0.746 |

**Table S10.** PAMPA test set performances of various machine learning regressors trained on SMILES embeddings from pre-trained language models (MoLFormer XL, PubChem10M SMILES BPE, and ChemBERTa-zinc-base-v1), fine-tuned on the PAMPA train dataset. Evaluation metrics include MSE, MAE, RMSE, R², PCC, and SCC.

| **Chemical Language models** | **Metrics** | **LGBM** | **Decision Tree** | **Random Forest** | **Gradient Boosting** | **AdaBoost** | **XGBoost** | **Extra Trees** | **Linear Regression** | **KNN** | **SVR** | **MLP** |
| --- | --- | --- | --- | --- | --- | --- | --- | --- | --- | --- | --- | --- |
| **MolFormer_XL_both-10pct** | **MSE** | 0.206 | 0.239 | 0.208 | 0.211 | 0.233 | 0.208 | **0.206** | 0.216 | 0.217 | 0.208 | 0.271 |
|  | **MAE** | 0.325 | 0.347 | 0.325 | 0.330 | 0.366 | 0.326 | **0.322** | 0.336 | 0.333 | 0.322 | 0.373 |
|  | **RMSE** | 0.454 | 0.489 | 0.456 | 0.459 | 0.483 | 0.457 | **0.454** | 0.464 | 0.465 | 0.456 | 0.520 |
|  | **R^2^** | 0.676 | 0.624 | 0.672 | 0.669 | 0.634 | 0.672 | **0.676** | 0.661 | 0.659 | 0.674 | 0.574 |
|  | **PCC** | 0.822 | 0.795 | 0.820 | 0.818 | 0.802 | 0.821 | **0.822** | 0.814 | 0.814 | 0.821 | 0.772 |
|  | **SCC** | 0.804 | 0.775 | 0.803 | 0.798 | 0.779 | 0.804 | **0.806** | 0.791 | 0.801 | 0.807 | 0.746 |
| **PubChem10M_SMILES_BPE_450K** | **MSE** | 0.266 | 0.286 | 0.264 | 0.269 | 0.269 | 0.263 | 0.262 | 0.267 | 0.268 | **0.262** | 0.314 |
|  | **MAE** | 0.365 | 0.385 | 0.363 | 0.367 | 0.379 | 0.365 | 0.362 | 0.365 | 0.365 | **0.361** | 0.402 |
|  | **RMSE** | 0.516 | 0.535 | 0.513 | 0.519 | 0.518 | 0.513 | 0.512 | 0.517 | 0.518 | **0.512** | 0.560 |
|  | **R^2^** | 0.582 | 0.550 | 0.586 | 0.577 | 0.577 | 0.586 | 0.588 | 0.580 | 0.578 | **0.588** | 0.506 |
|  | **PCC** | 0.769 | 0.754 | 0.771 | 0.766 | 0.763 | 0.772 | 0.772 | 0.770 | 0.770 | **0.772** | 0.752 |
|  | **SCC** | 0.751 | 0.728 | 0.754 | 0.748 | 0.742 | 0.754 | 0.754 | 0.753 | 0.754 | **0.757** | 0.738 |
| **ChemBERTa-zinc-base-v1** | **MSE** | 0.270 | 0.287 | 0.272 | 0.275 | 0.278 | 0.273 | 0.274 | 0.293 | 0.281 | **0.270** | 0.304 |
|  | **MAE** | 0.364 | 0.382 | 0.365 | 0.369 | 0.389 | 0.366 | 0.365 | 0.377 | 0.372 | **0.366** | 0.386 |
|  | **RMSE** | 0.520 | 0.536 | 0.521 | 0.524 | 0.528 | 0.522 | 0.523 | 0.541 | 0.530 | **0.520** | 0.552 |
|  | **R^2^** | 0.575 | 0.549 | 0.573 | 0.568 | 0.562 | 0.571 | 0.570 | 0.539 | 0.559 | **0.576** | 0.521 |
|  | **PCC** | 0.764 | 0.751 | 0.762 | 0.759 | 0.753 | 0.762 | 0.762 | 0.745 | 0.757 | **0.764** | 0.742 |
|  | **SCC** | 0.751 | 0.733 | 0.749 | 0.745 | 0.737 | 0.749 | 0.747 | 0.739 | 0.749 | **0.754** | 0.732 |

**Table S11.** Test set performance metrics of models trained on top 50 combined features (including atomic features, molecular fingerprints, descriptors, and molecular embeddings) for PAMPA permeability prediction.

|  | **MSE** | **MAE** | **RMSE** | **R^2^** | **PCC** | **SCC** |
| --- | --- | --- | --- | --- | --- | --- |
| **LGBM** | 0.210 | 0.329 | 0.458 | 0.670 | 0.819 | 0.798 |
| **Decision Trees** | 0.242 | 0.350 | 0.492 | 0.620 | 0.792 | 0.771 |
| **Random Forest** | 0.209 | 0.326 | 0.457 | 0.671 | 0.820 | 0.801 |
| **Gradient Boosting** | 0.214 | 0.332 | 0.462 | 0.664 | 0.816 | 0.794 |
| **AdaBoost** | 0.240 | 0.369 | 0.489 | 0.623 | 0.794 | 0.773 |
| **XGBoost** | **0.207** | **0.325** | **0.455** | **0.674** | **0.822** | **0.802** |
| **Extra Trees** | 0.209 | 0.325 | 0.457 | 0.671 | 0.820 | 0.802 |
| **Linear Regression** | 0.216 | 0.335 | 0.465 | 0.660 | 0.813 | 0.789 |
| **KNN** | 0.223 | 0.337 | 0.472 | 0.650 | 0.809 | 0.789 |
| **SVR** | 0.213 | 0.327 | 0.462 | 0.664 | 0.816 | 0.798 |
| **MLP** | 0.224 | 0.343 | 0.474 | 0.647 | 0.807 | 0.775 |

**Table S12.** Predictive performance of machine learning models on the Caco2 test dataset using atomic features alone versus a combination of atomic features and monomeric composition. Evaluation metrics include MSE, MAE, RMSE, R², PCC, and SCC.

|  | **Atomic features** | | | | | | **Atomic features + monomeric composition** | | | | | |
| --- | --- | --- | --- | --- | --- | --- | --- | --- | --- | --- | --- | --- |
| **Models** | **MSE** | **MAE** | **RMSE** | **R^2^** | **PCC** | **SCC** | **MSE** | **MAE** | **RMSE** | **R^2^** | **PCC** | **SCC** |
| **LGBM** | 0.186 | 0.332 | 0.432 | 0.701 | 0.838 | 0.805 | 0.160 | 0.308 | 0.400 | 0.743 | 0.863 | 0.849 |
| **Decision Trees** | 0.223 | 0.350 | 0.472 | 0.642 | 0.808 | 0.777 | 0.184 | 0.316 | 0.429 | 0.705 | 0.841 | 0.811 |
| **Random Forest** | 0.184 | 0.327 | 0.428 | 0.706 | 0.840 | 0.810 | 0.146 | 0.288 | 0.381 | 0.767 | 0.877 | 0.859 |
| **Gradient Boosting** | 0.199 | 0.349 | 0.446 | 0.681 | 0.828 | 0.792 | 0.173 | 0.324 | 0.416 | 0.722 | 0.854 | 0.827 |
| **AdaBoost** | 0.294 | 0.459 | 0.542 | 0.528 | 0.748 | 0.686 | 0.267 | 0.435 | 0.516 | 0.572 | 0.784 | 0.727 |
| **XGBoost** | 0.187 | 0.325 | 0.432 | 0.700 | 0.838 | 0.814 | **0.138** | **0.279** | **0.372** | **0.778** | **0.883** | **0.866** |
| **Extra Trees** | **0.169** | **0.317** | **0.411** | **0.729** | **0.854** | **0.829** | 0.156 | 0.287 | 0.395 | 0.750 | 0.866 | 0.836 |
| **Linear Regression** | 0.384 | 0.513 | 0.620 | 0.384 | 0.625 | 0.528 | 0.404 | 0.429 | 0.636 | 0.352 | 0.702 | 0.716 |
| **KNN** | 0.209 | 0.359 | 0.458 | 0.664 | 0.815 | 0.775 | 0.278 | 0.399 | 0.527 | 0.554 | 0.746 | 0.712 |
| **SVR** | 0.225 | 0.373 | 0.475 | 0.639 | 0.800 | 0.751 | 0.206 | 0.349 | 0.453 | 0.670 | 0.821 | 0.801 |
| **MLP** | 0.412 | 0.494 | 0.642 | 0.339 | 0.637 | 0.616 | 0.626 | 0.544 | 0.791 | -0.004 | 0.657 | 0.703 |

**Table S13.** Performance comparison of machine learning models on the Caco2 test dataset using monomer composition and their natural amino acid analogs as input features. Test set performance metrics include MSE, MAE, RMSE, R², PCC, and SCC for evaluating permeability prediction.

|  | **Monomer Composition** | | | | | | **Natural analog monomeric Composition** | | | | | |
| --- | --- | --- | --- | --- | --- | --- | --- | --- | --- | --- | --- | --- |
| **Models** | **MSE** | **MAE** | **RMSE** | **R^2^** | **PCC** | **SCC** | **MSE** | **MAE** | **RMSE** | **R^2^** | **PCC** | **SCC** |
| **Extra Trees** | 0.189 | 0.322 | 0.435 | 0.697 | 0.835 | 0.812 | **0.229** | **0.352** | **0.478** | **0.633** | **0.797** | **0.757** |
| **LGBM** | 0.219 | 0.357 | 0.468 | 0.649 | 0.806 | 0.792 | 0.260 | 0.393 | 0.510 | 0.583 | 0.763 | 0.721 |
| **XGBoost** | **0.181** | **0.320** | **0.425** | **0.710** | **0.843** | **0.826** | 0.249 | 0.378 | 0.499 | 0.601 | 0.780 | 0.739 |
| **Decision Tree** | 0.224 | 0.351 | 0.473 | 0.641 | 0.804 | 0.780 | 0.255 | 0.369 | 0.505 | 0.591 | 0.777 | 0.731 |
| **Random Forest** | 0.183 | 0.318 | 0.427 | 0.707 | 0.842 | 0.826 | 0.234 | 0.362 | 0.484 | 0.624 | 0.791 | 0.746 |
| **Gradient Boosting** | 0.222 | 0.363 | 0.471 | 0.644 | 0.812 | 0.777 | 0.249 | 0.402 | 0.499 | 0.601 | 0.780 | 0.745 |
| **AdaBoost** | 0.385 | 0.531 | 0.621 | 0.382 | 0.642 | 0.603 | 0.382 | 0.529 | 0.618 | 0.386 | 0.631 | 0.614 |
| **SVR** | 0.270 | 0.383 | 0.519 | 0.567 | 0.754 | 0.742 | 0.294 | 0.423 | 0.543 | 0.528 | 0.727 | 0.702 |
| **Linear Regression** | 0.310 | 0.410 | 0.557 | 0.503 | 0.736 | 0.741 | 0.466 | 0.573 | 0.683 | 0.252 | 0.503 | 0.441 |
| **KNN** | 0.369 | 0.447 | 0.607 | 0.408 | 0.656 | 0.637 | 0.314 | 0.422 | 0.560 | 0.496 | 0.713 | 0.685 |
| **MLP** | 0.781 | 0.615 | 0.884 | -0.253 | 0.585 | 0.646 | 0.714 | 0.648 | 0.845 | -0.145 | 0.418 | 0.411 |

**Table S14.** Predictive performances of best-performing regressors trained on different molecular fingerprint types for the Caco2 test dataset. Test set performance metrics reported include MSE, MAE, RMSE, R², PCC, and SCC.

| **Fingerprint Type** | **Model** | **MSE** | **MAE** | **RMSE** | **R^2^** | **PCC** | **SCC** |
| --- | --- | --- | --- | --- | --- | --- | --- |
| **All fingerprints** | XGBoost | **0.134** | **0.280** | **0.366** | **0.785** | **0.887** | **0.870** |
| **AtomPairs2D** | Gradient Boosting | 0.383 | 0.510 | 0.619 | 0.386 | 0.621 | 0.507 |
| **Count AtomPairs2D** | Extra Trees | 0.147 | 0.295 | 0.383 | 0.764 | 0.875 | 0.859 |
| **Count KlekotaRoth** | Extra trees | 0.165 | 0.302 | 0.406 | 0.735 | 0.858 | 0.836 |
| **Count Morgan** | XGBoost | **0.144** | **0.284** | **0.380** | **0.769** | **0.878** | **0.863** |
| **Count Substructure** | XGBoost | 0.159 | 0.298 | 0.399 | 0.745 | 0.864 | 0.850 |
| **EState** | Extra trees | 0.386 | 0.504 | 0.621 | 0.381 | 0.618 | 0.555 |
| **Extended** | XGBoost | 0.254 | 0.387 | 0.504 | 0.592 | 0.771 | 0.739 |
| **Fingerprinter** | XGBoost | 0.265 | 0.395 | 0.515 | 0.575 | 0.762 | 0.718 |
| **GraphOnly** | Random Forest | 0.289 | 0.418 | 0.538 | 0.536 | 0.733 | 0.682 |
| **KlekotaRoth** | XGBoost | 0.187 | 0.331 | 0.432 | 0.700 | 0.837 | 0.795 |
| **MACCS** | SVR | 0.252 | 0.399 | 0.502 | 0.596 | 0.773 | 0.723 |
| **Morgan** | Random Forest | 0.172 | 0.317 | 0.415 | 0.723 | 0.854 | 0.828 |
| **PubChem** | SVR | 0.287 | 0.422 | 0.536 | 0.539 | 0.734 | 0.675 |
| **Substructure** | Random Forest | 0.362 | 0.486 | 0.602 | 0.419 | 0.649 | 0.606 |

**Table S15.** Comparative test set performances of different pre-trained molecular language models, MoLFormer XL both-10pct, PubChem10M SMILES BPE 450K, and ChemBERTa-zinc-base-v1 fine-tuned on the Caco-2 training set. Evaluation on the test set was conducted using standard regression metrics: MSE, MAE, RMSE, R², PCC, and SCC.

| **Metrics** | **MolFormer_XL_both-10pct** | **PubChem10M_SMILES_BPE_450K** | **ChemBERTa-zinc-base-v1** |
| --- | --- | --- | --- |
| **MSE** | **0.296** | 0.349 | 0.322 |
| **MAE** | **0.435** | 0.442 | 0.445 |
| **RMSE** | **0.544** | 0.591 | 0.568 |
| **R^2^** | **0.526** | 0.440 | 0.483 |
| **PCC** | **0.798** | 0.760 | 0.719 |
| **SCC** | **0.765** | 0.730 | 0.681 |

**Table S16.** Caco2 test set performances of various machine learning regressors trained on SMILES embeddings from pre-trained language models (MoLFormer XL, PubChem10M SMILES BPE, and ChemBERTa-zinc-base-v1), fine-tuned on the Caco2 train dataset.

| **Chemical Language Model** | **Metrics** | **LGBM** | **Decision Tree** | **Random Forest** | **Gradient Boosting** | **AdaBoost** | **XGBoost** | **Extra Trees** | **Linear Regression** | **KNN** | **SVR** | **MLP** |
| --- | --- | --- | --- | --- | --- | --- | --- | --- | --- | --- | --- | --- |
| **MolFormer_XL_both-10pct** | **MSE** | 0.187 | 0.229 | 0.190 | 0.196 | 0.215 | 0.193 | **0.182** | 1.478 | 0.185 | 0.183 | 0.412 |
|  | **MAE** | 0.323 | 0.360 | 0.329 | 0.332 | 0.364 | 0.330 | **0.320** | 0.932 | 0.326 | 0.325 | 0.481 |
|  | **RMSE** | 0.433 | 0.479 | 0.436 | 0.443 | 0.463 | 0.439 | **0.426** | 1.216 | 0.430 | 0.428 | 0.642 |
|  | **R^2^** | 0.699 | 0.632 | 0.694 | 0.685 | 0.656 | 0.690 | **0.708** | -1.372 | 0.703 | 0.706 | 0.338 |
|  | **PCC** | 0.836 | 0.797 | 0.834 | 0.828 | 0.811 | 0.831 | **0.842** | 0.253 | 0.839 | 0.842 | 0.690 |
|  | **SCC** | 0.789 | 0.763 | 0.789 | 0.779 | 0.759 | 0.790 | **0.799** | 0.272 | 0.799 | 0.807 | 0.664 |
| **PubChem10M_SMILES_BPE_450K** | **MSE** | 0.252 | 0.261 | 0.249 | 0.252 | 0.259 | 0.253 | 0.250 | 0.947 | **0.240** | 0.251 | 0.426 |
|  | **MAE** | 0.373 | 0.383 | 0.371 | 0.374 | 0.384 | 0.374 | 0.371 | 0.744 | **0.357** | 0.374 | 0.506 |
|  | **RMSE** | 0.502 | 0.511 | 0.499 | 0.502 | 0.509 | 0.503 | 0.500 | 0.973 | **0.490** | 0.501 | 0.653 |
|  | **R^2^** | 0.596 | 0.582 | 0.600 | 0.595 | 0.585 | 0.594 | 0.600 | -0.520 | **0.615** | 0.597 | 0.317 |
|  | **PCC** | 0.779 | 0.771 | 0.780 | 0.779 | 0.770 | 0.778 | 0.780 | 0.443 | **0.793** | 0.776 | 0.621 |
|  | **SCC** | 0.739 | 0.725 | 0.739 | 0.737 | 0.725 | 0.739 | 0.739 | 0.422 | **0.760** | 0.738 | 0.556 |
| **ChemBERTa-zinc-base-v1** | **MSE** | 0.293 | 0.309 | 0.295 | 0.296 | 0.297 | 0.288 | 0.289 | 1.280 | **0.286** | 0.280 | 0.688 |
|  | **MAE** | 0.407 | 0.419 | 0.410 | 0.409 | 0.418 | 0.407 | 0.404 | 0.863 | **0.393** | 0.398 | 0.631 |
|  | **RMSE** | 0.541 | 0.556 | 0.543 | 0.544 | 0.545 | 0.537 | 0.537 | 1.132 | **0.535** | 0.529 | 0.829 |
|  | **R^2^** | 0.530 | 0.504 | 0.527 | 0.524 | 0.524 | 0.537 | 0.537 | -1.055 | **0.541** | 0.551 | -0.104 |
|  | **PCC** | 0.739 | 0.725 | 0.735 | 0.736 | 0.732 | 0.743 | 0.741 | 0.264 | **0.748** | 0.749 | 0.629 |
|  | **SCC** | 0.694 | 0.681 | 0.689 | 0.693 | 0.689 | 0.698 | 0.695 | 0.278 | **0.716** | 0.712 | 0.582 |

**Table S17.** Comparative test set performance metrics of models trained on top 50 features (including atomic features, molecular fingerprints, descriptors, and molecular embeddings) for Caco-2 permeability prediction. Test set performance metrics reported include MSE, MAE, RMSE, R², PCC, and SCC.

|  | MSE | MAE | RMSE | R2 | PCC | SCC |
| --- | --- | --- | --- | --- | --- | --- |
| LGBM | 0.194 | 0.333 | 0.440 | 0.689 | 0.831 | 0.786 |
| Decision Trees | 0.223 | 0.350 | 0.472 | 0.642 | 0.804 | 0.757 |
| Random Forest | 0.196 | 0.334 | 0.442 | 0.686 | 0.828 | 0.784 |
| Gradient Boosting | 0.198 | 0.340 | 0.445 | 0.683 | 0.827 | 0.780 |
| AdaBoost | 0.220 | 0.371 | 0.469 | 0.647 | 0.806 | 0.752 |
| XGBoost | 0.199 | 0.340 | 0.446 | 0.681 | 0.826 | 0.783 |
| Extra Trees | 0.186 | 0.326 | 0.431 | 0.702 | 0.838 | 0.792 |
| Linear Regression | 0.219 | 0.361 | 0.468 | 0.649 | 0.809 | 0.776 |
| KNN | 0.184 | 0.327 | 0.429 | 0.705 | 0.841 | 0.794 |
| SVR | **0.184** | **0.326** | **0.429** | **0.705** | **0.841** | **0.803** |
| MLP | 0.463 | 0.513 | 0.681 | 0.257 | 0.652 | 0.641 |

**Supplementary Figures**

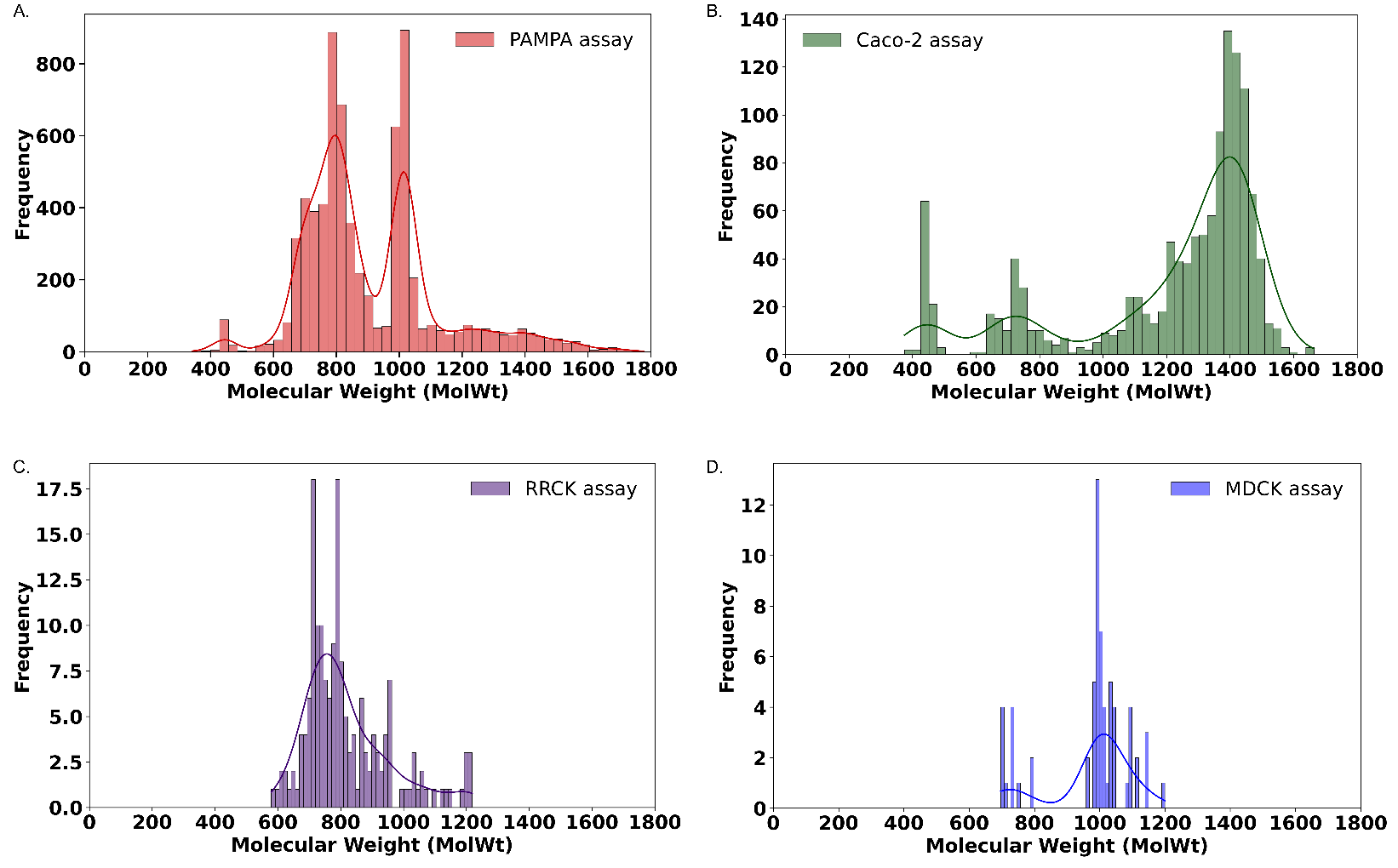

**Figure S1.** Distribution of molecular weights of peptides across different permeability assay datasets. The histograms illustrate the dispersion of molecular weights (in Daltons) measured in: (A) PAMPA assay (B) Caco-2 assay, (C) RRCK assay, and (D) MDCK assay. Each panel includes a kernel density estimation (KDE) curve to visualize the distribution trends.

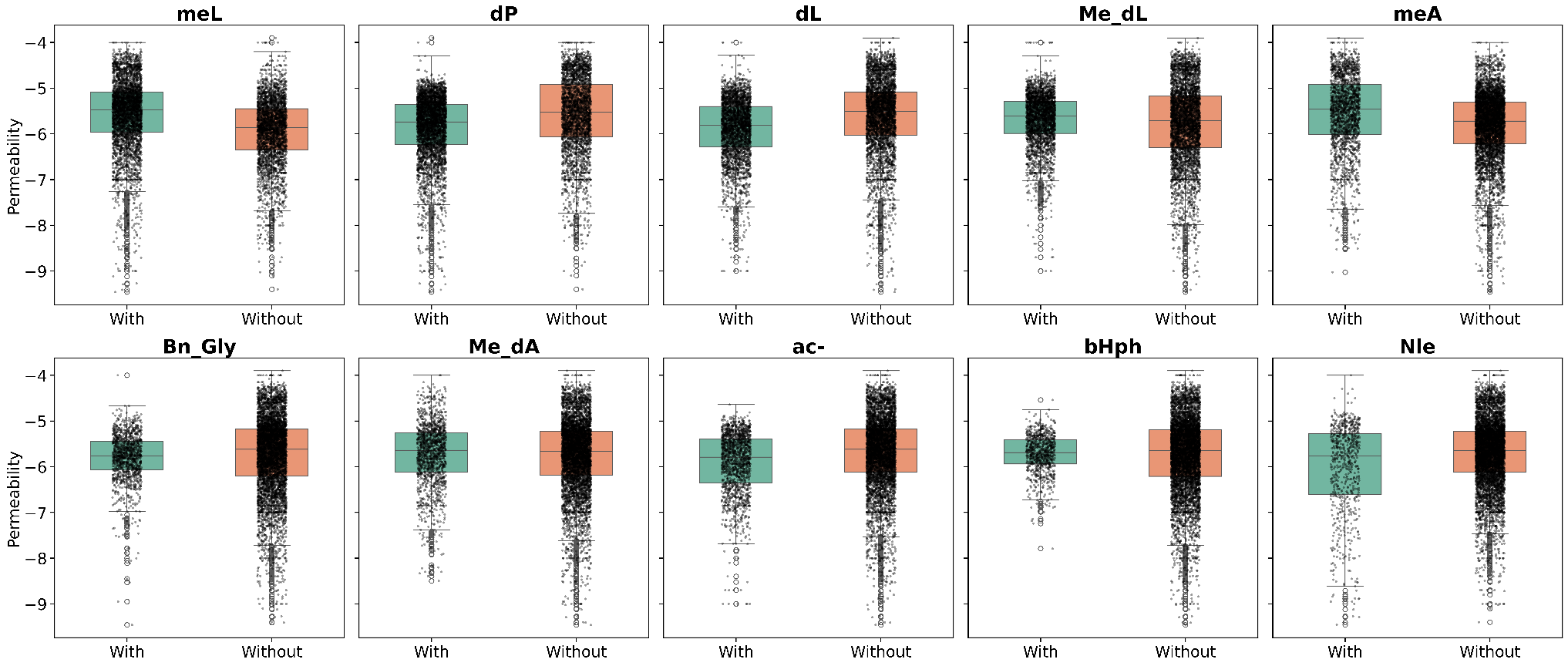

**Figure S2.** Effect of the ten most abundant non-canonical monomers on peptide permeability. Boxplots show permeability distributions of peptides containing (“with”) or lacking (“without”) each monomer in PAMPA datasets, using -6 as the threshold for permeable versus non-permeable peptides. Peptides having permeability higher than -6 are considered permeable.

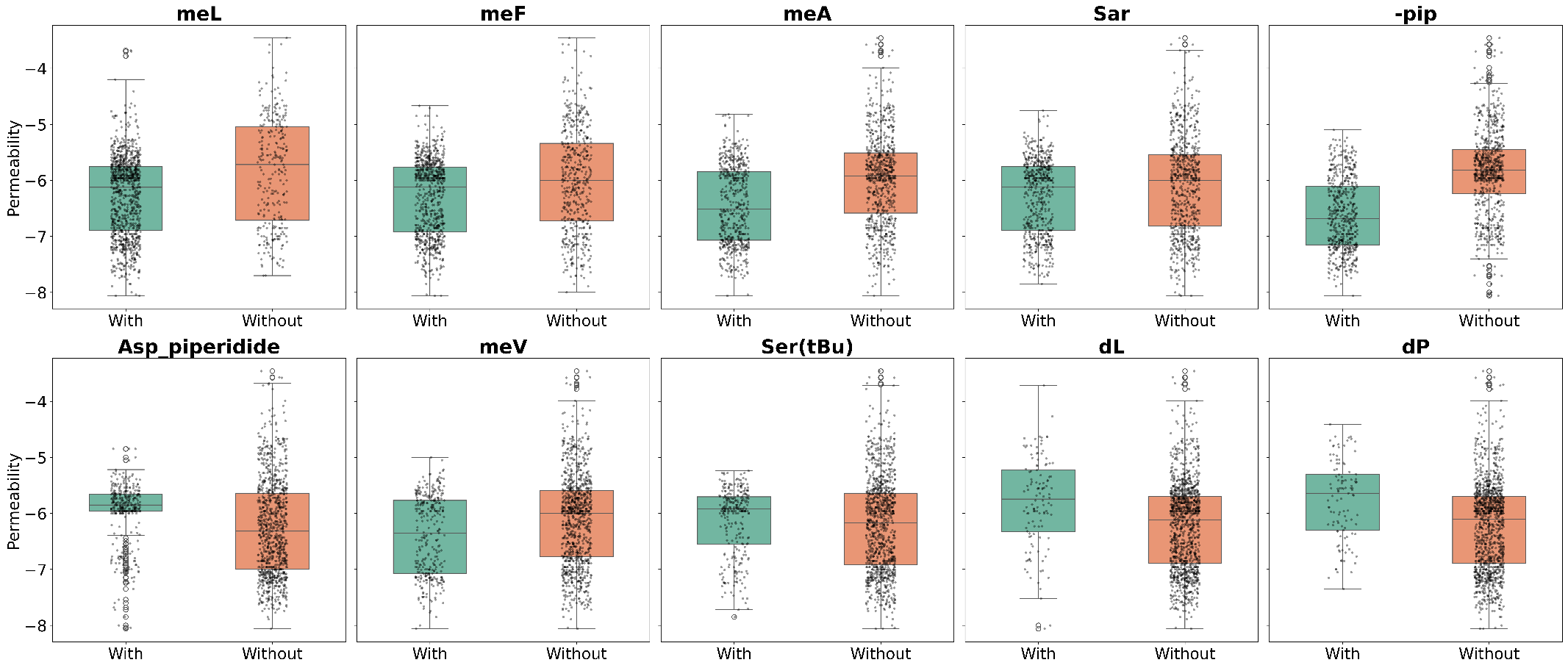

**Figure S3.** Effect of the ten most abundant non-canonical monomers on peptide permeability. Boxplots show permeability distributions of peptides containing (“with”) or lacking (“without”) each monomer in Caco-2 datasets, using -6 as the threshold for permeable versus non-permeable peptides. Peptides having permeability higher than -6 are considered permeable.

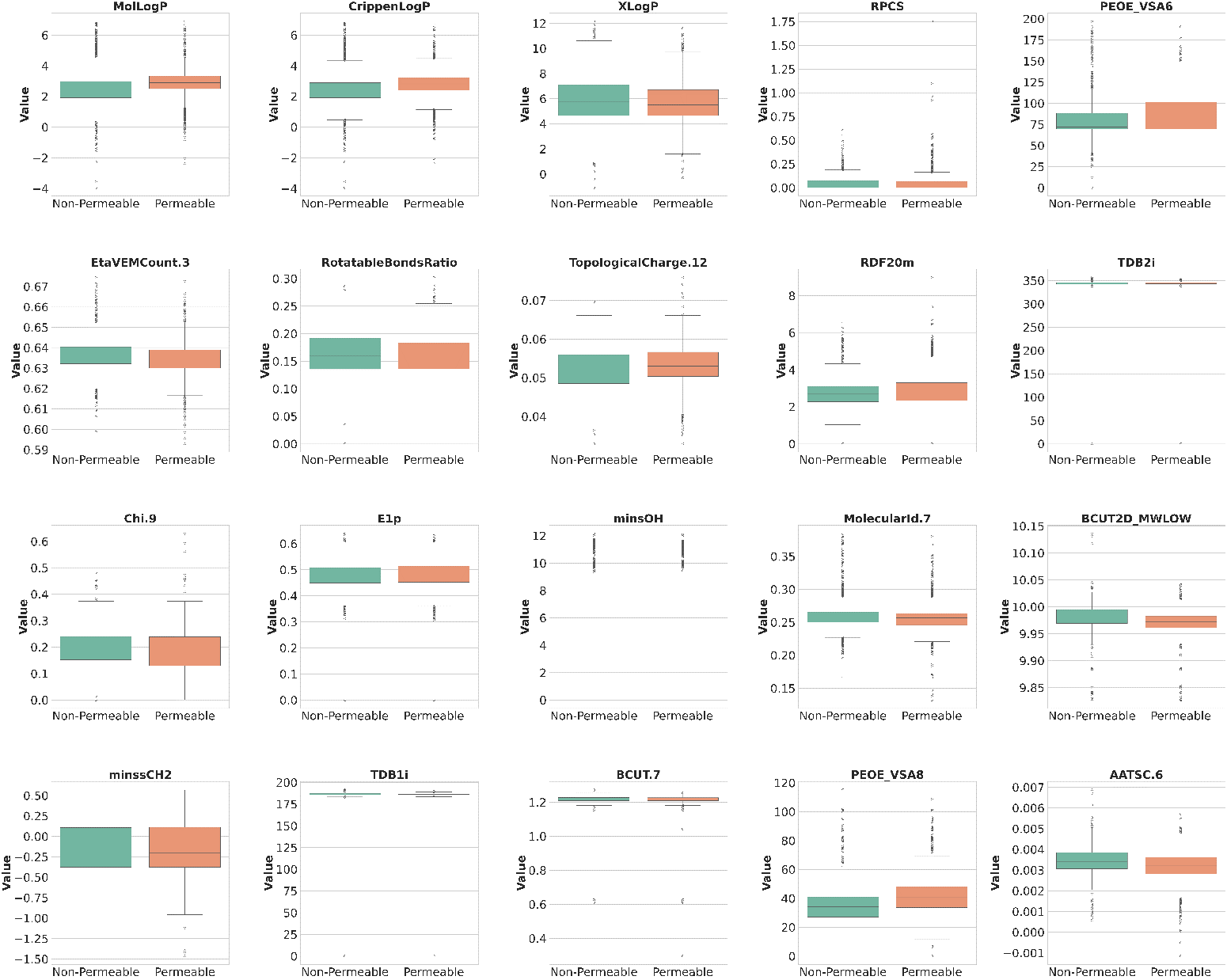

**Figure S4.** Boxplots illustrating the distribution of the top 20 most important molecular descriptors for peptides in the PAMPA training dataset. Each subplot compares descriptor values between permeable and non-permeable peptide classes. These descriptors capture key physicochemical and topological properties influencing passive membrane permeability.

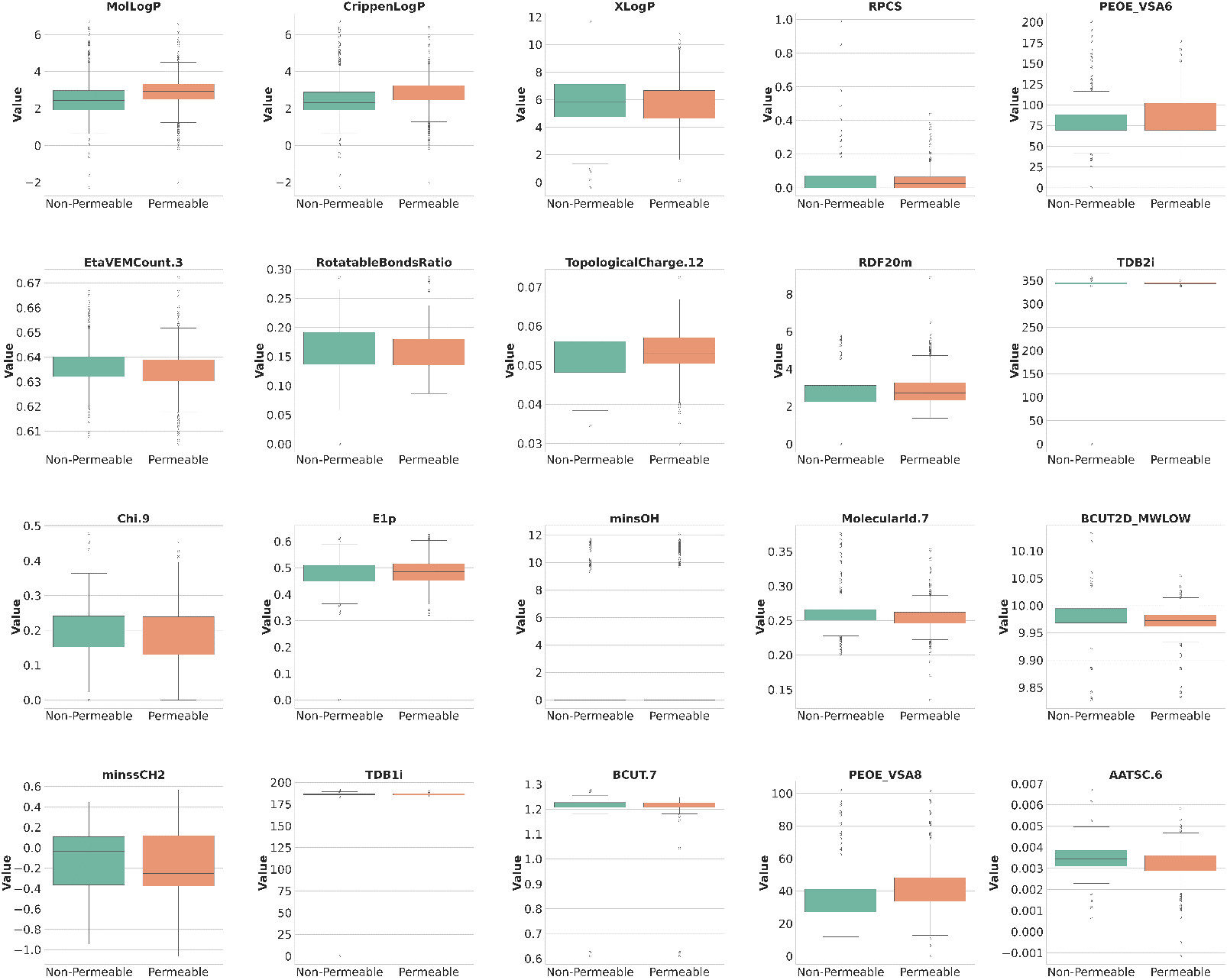

**Figure S5.** Boxplots illustrating the distribution of the top 20 most important molecular descriptors for peptides in the PAMPA test set. Each subplot compares descriptor values between permeable and non-permeable peptide classes. These descriptors capture key physicochemical and topological properties influencing passive membrane permeability.

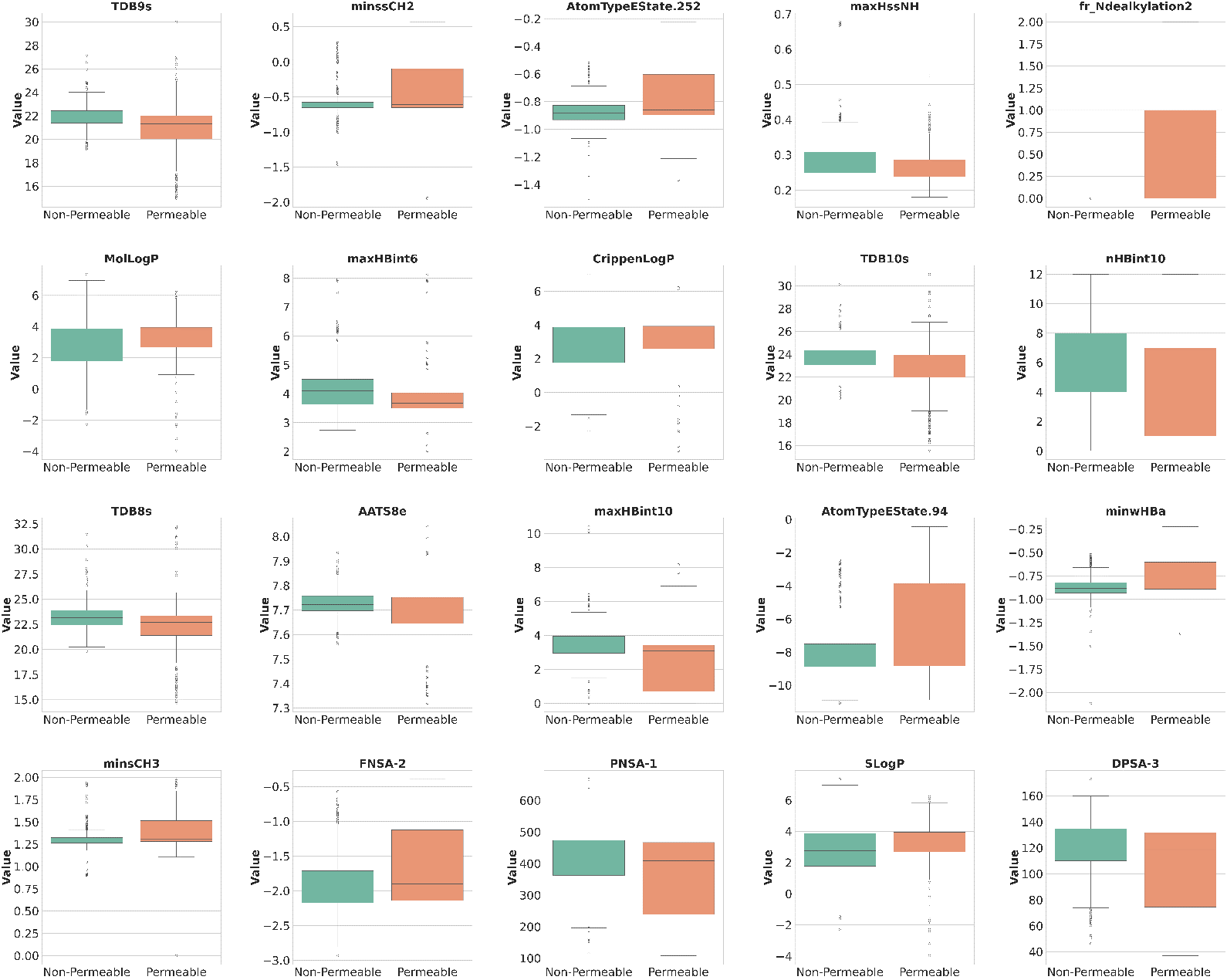

**Figure S6.** Boxplots showing the distribution of the top 20 most important molecular descriptors (features) for peptides in the Caco2 training dataset. Each subplot compares the descriptor values between permeable and non-permeable peptide classes. The descriptors reflect physicochemical and topological properties contributing to passive membrane permeability classification.

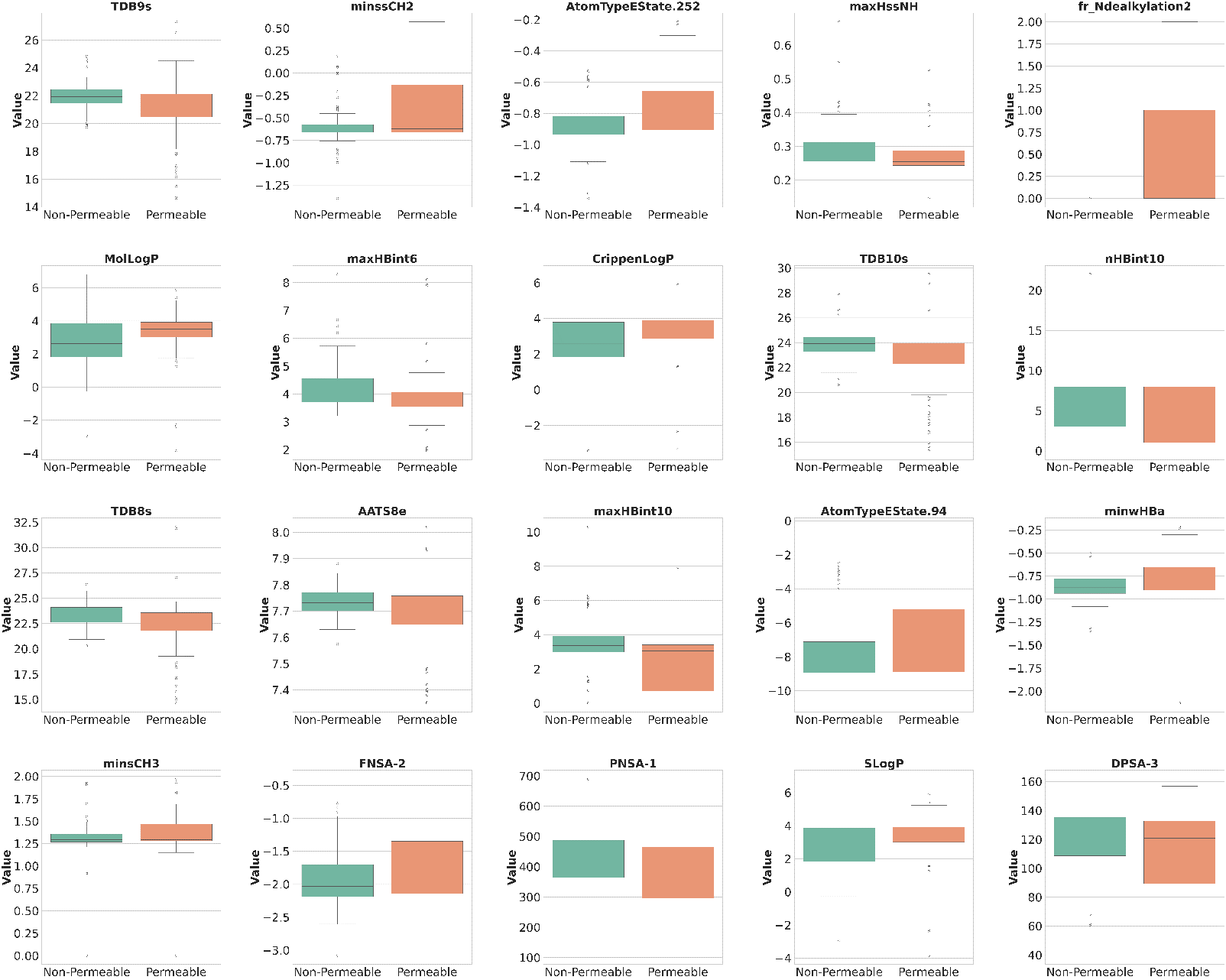

**Figure S7.** Boxplots showing the distribution of the top 20 most important molecular descriptors (features) for peptides in the Caco2 test dataset. Each subplot compares the descriptor values between permeable and non-permeable peptide classes. The descriptors reflect physicochemical and topological properties contributing to passive membrane permeability classification.

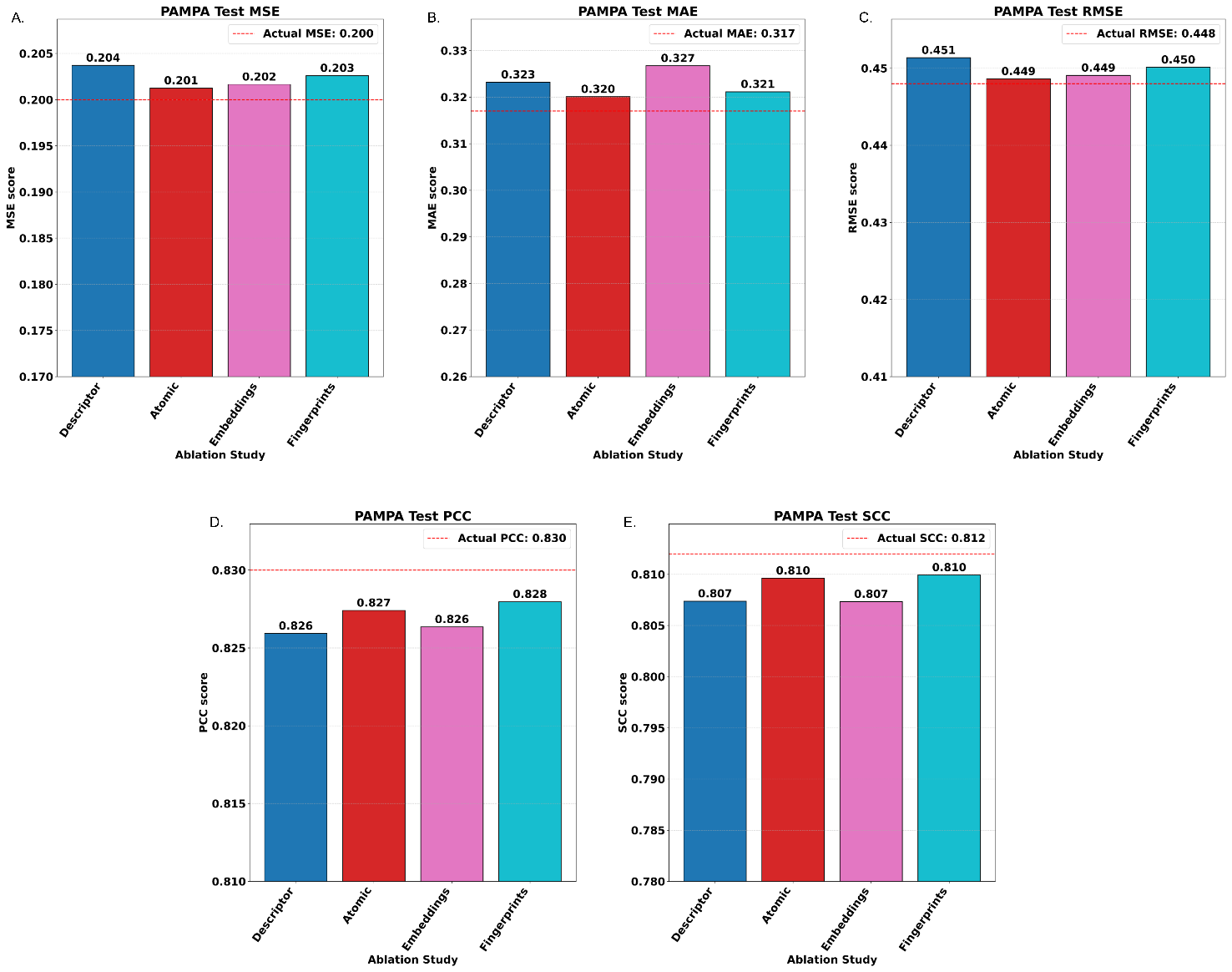

**Figure S8.** Ablation study on stacked ensemble architecture used to predict permeability of peptides for PAMPA data. Each bar plot (A-E) shows the effect of systematically excluding models trained on one feature type on the following performance metrics: (A) MSE, (B) MAE, (C) RMSE, (D) PCC, and (E) SCC. The dashed red line in each plot represents the performance of the full model utilizing all feature types, serving as a reference benchmark.

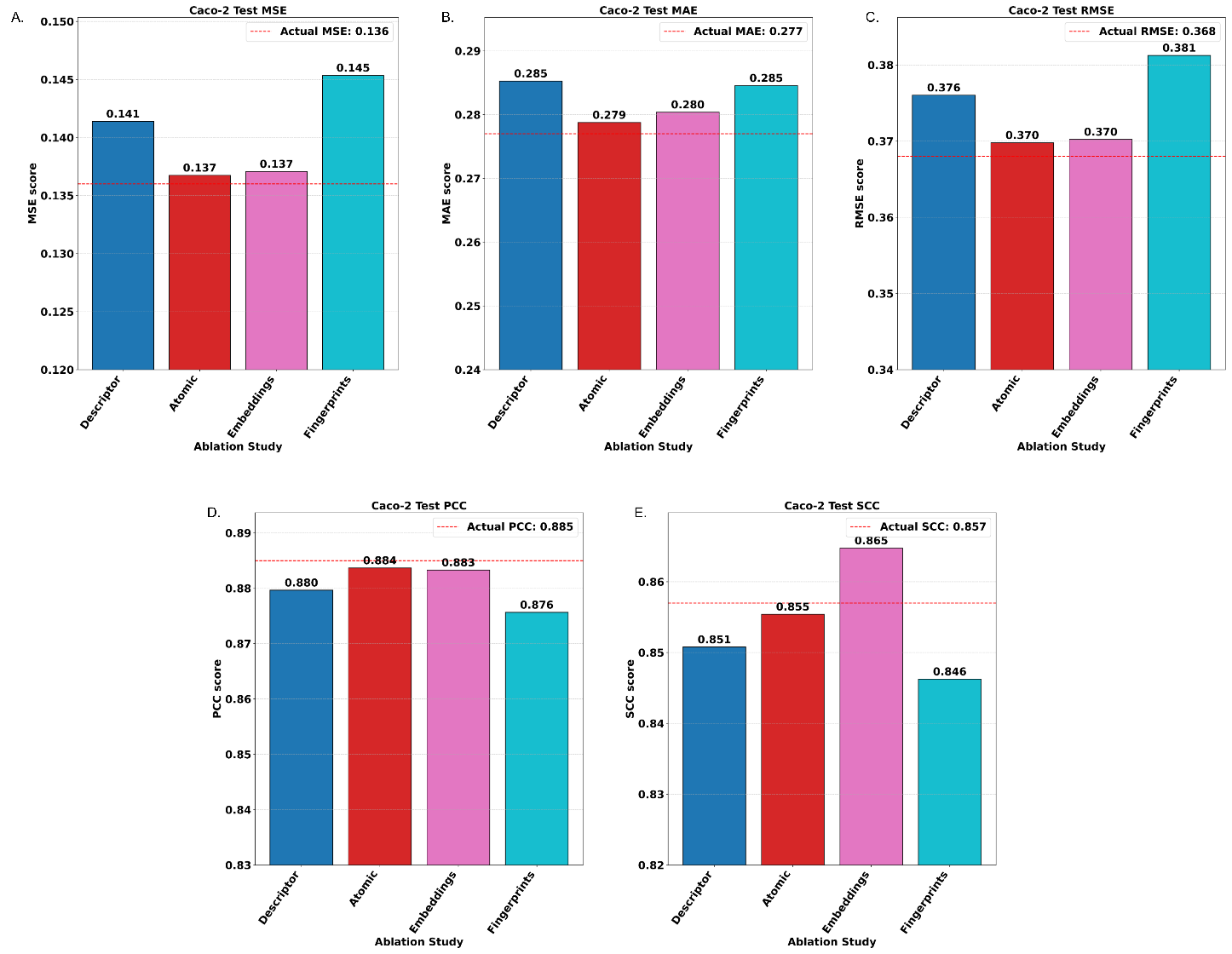

**Figure S9.** Ablation study on the stacked ensemble architecture used to predict the permeability of peptides in Caco-2 assay data. Each bar plot (A-E) reflects the effect of systematically excluding models trained on one feature type on the following performance metrics: (A) MSE, (B) MAE, (C) RMSE, (D) PCC, and (E) SCC. The dashed red line in each plot represents the performance of the full model utilizing all feature types, serving as a reference benchmark.

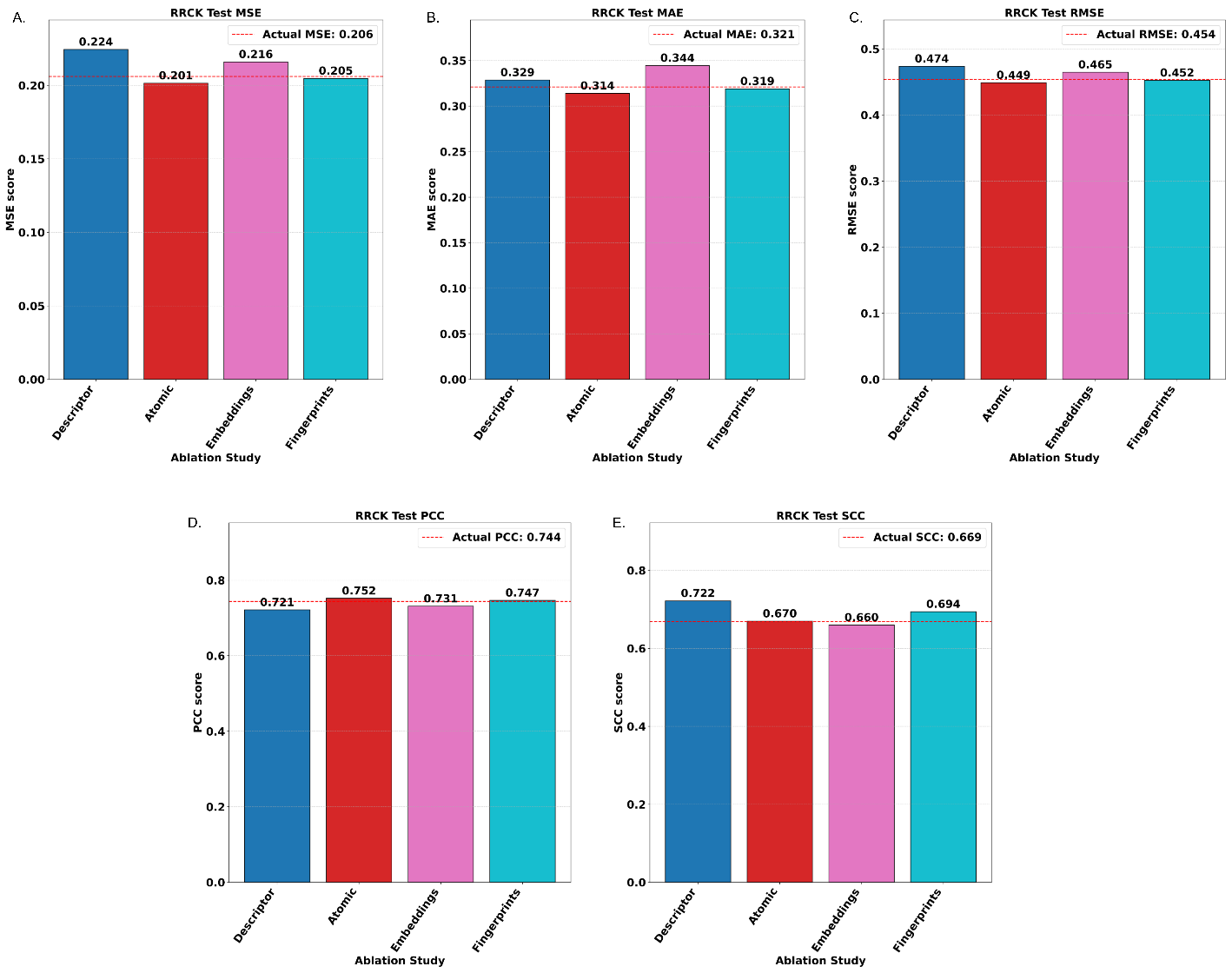

**Figure S10.** Ablation study on stacked ensemble architecture used to predict the permeability of peptides in RRCK assay data. Each bar plot (A-E) presents the effect of removing models trained on one feature type on the following performance metrics: (A) MSE, (B) MAE, (C) RMSE, (D) PCC, and (E) SCC. The dashed red line in each plot represents the performance of the full model utilizing all feature types, serving as a reference benchmark.

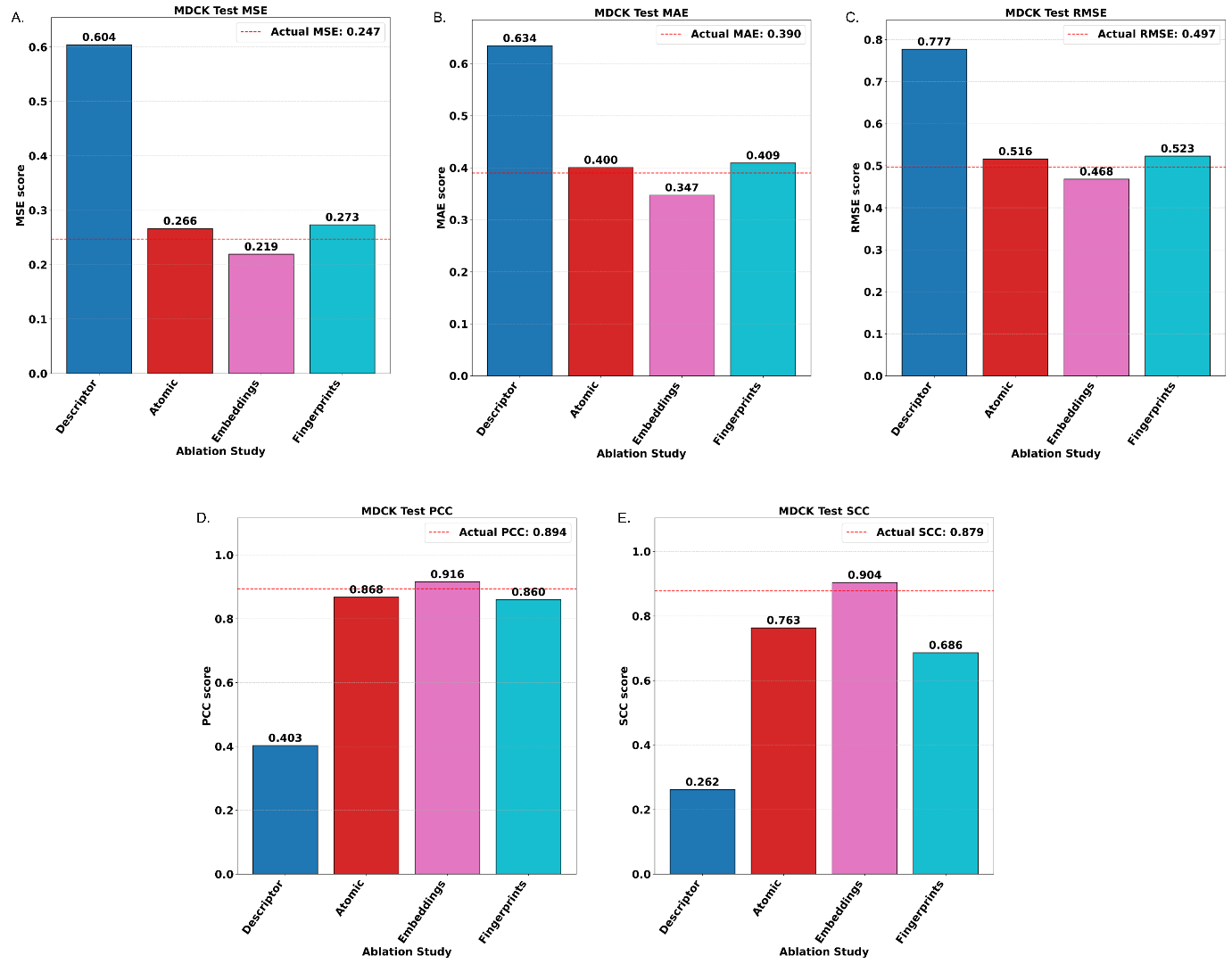

**Figure S11.** Ablation study on stacked ensemble architecture used to predict the permeability of peptides in MDCK assay data. Each bar plot (A-E) presents the effect of removing models trained on one feature type on the following performance metrics: (A) MSE, (B) MAE, (C) RMSE, (D) PCC, and (E) SCC. The dashed red line in each plot represents the performance of the full model utilizing all feature types, serving as a reference benchmark.
